# Bacterial retrons encode tripartite toxin/antitoxin systems

**DOI:** 10.1101/2020.06.22.160168

**Authors:** Jacob Bobonis, André Mateus, Birgit Pfalz, Sarela Garcia-Santamarina, Marco Galardini, Callie Kobayashi, Frank Stein, Mikhail M. Savitski, Johanna R. Elfenbein, Helene Andrews-Polymenis, Athanasios Typas

## Abstract

Retrons are genetic retroelements, commonly found in bacterial genomes and recently repurposed as genome editing tools. Their encoded reverse transcriptase (RT) produces a multi-copy single-stranded DNA (msDNA). Despite our understanding of their complex biosynthesis, the function of msDNAs and therefore, the physiological role of retrons has remained elusive. We establish that the retron-Sen2 in *Salmonella* Typhimurium encodes a toxin, which we have renamed as RcaT (Retron cold-anaerobic Toxin). RcaT is activated when msDNA biosynthesis is perturbed and its toxicity is higher at ambient temperatures or during anaerobiosis. The RT and msDNA form together the antitoxin unit, with the RT binding RcaT, and the msDNA enabling the antitoxin activity. Using another *E. coli* retron, we establish that this toxin/antitoxin function is conserved, and that RT-toxin interactions are cognate. Altogether, retrons constitute a novel family of tripartite toxin/antitoxin systems.

## INTRODUCTION

Reverse transcriptases (RTs), thought initially to be unique in retroviruses, were first discovered three decades ago in bacteria, encoded in genetic elements named retrons ^1,2^. Retrons were originally identified because they produce multiple copies of satellite DNA molecules, called msDNA (multi-copy single-stranded DNA) ^3^. They are present in bacteria across the phylogenetic tree ^4^. The ability of retrons to produce high-quantities of single-stranded DNA *in situ* has been recently exploited for recombineering and other genome-editing approaches ^5–8^. Yet, the natural function of retrons has remained enigmatic.

In contrast to their function, the msDNA biogenesis pathway is well understood. Retrons contain a non-coding RNA gene (*msrmsd*), the RT, and often, accessory genes of unknown function ^9^ (Fig. 1A – here the retron-Sen2 [retron-ST85] depicted as example). RTs reverse transcribe their cognate msrmsd-RNA, utilising the msr-RNA as primer, and the msd-RNA as template ^10^ (Fig. 1B). While the msd-DNA is being reverse-transcribed, the msd-RNA template is degraded by ribonuclease H (RNAse H) ^10^ (Fig. 1A-B). The end product is usually a branched DNA/RNA hybrid, with the msd-DNA being covalently joined to the msr-RNA through a 2’-5’ phosphodiester bond made by the RT ^11^ (Fig. 1B). Some msDNAs (e.g., msDNA-Sen2) are pure unbranched DNA ^12,13^, with the DNA branch being separated from the RNA-branch by the housekeeping Exonuclease VII ^14^ (Exo VII; encoded by *xseA* and *xseB* genes; Fig. 1A). Exo VII cleaves four deoxyribonucleotides from the msDNA, separating it from its branched precursor (Fig. 1B). Both branched ^15^ and unbranched ^16^ msDNAs can remain in complex with their cognate RTs (Fig. 1B). The accessory proteins (e.g., STM14_4640 in retron-Sen2 ^17^, Fig. 1A), do not affect msDNA biosynthesis ^13,18^, and neither their sequence nor the position in retron elements are conserved.

**Figure 1.**
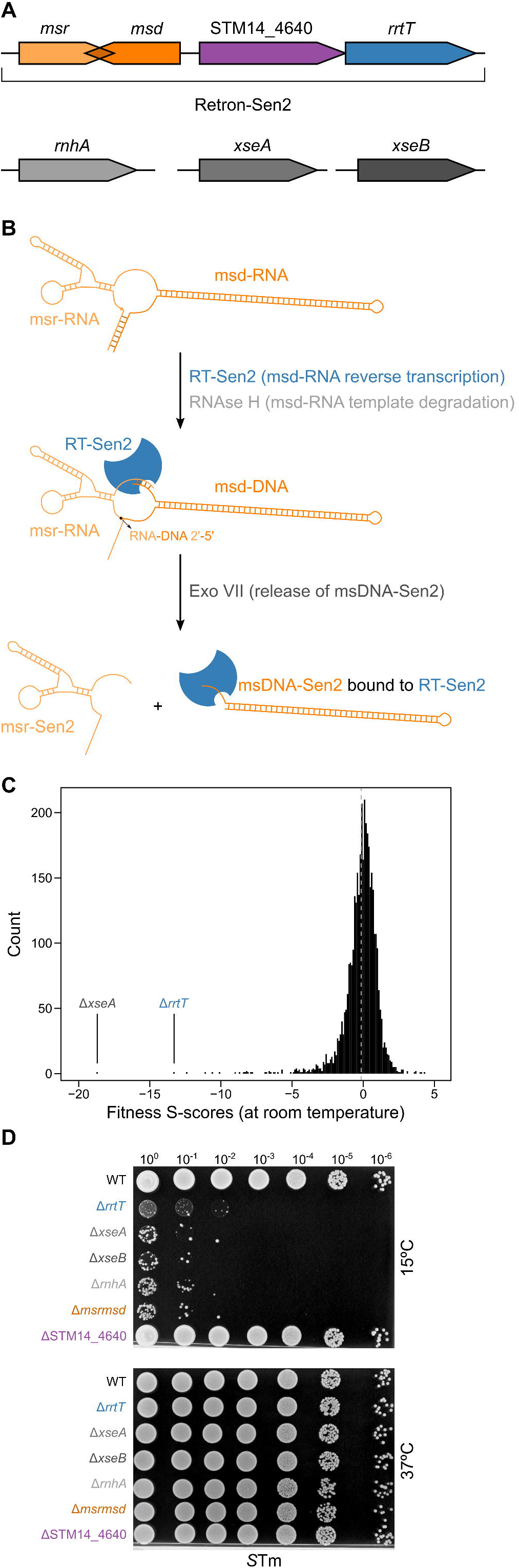
*S*Tm Retron-Sen2 deletion mutants are cold-sensitive. **(A)** Genes involved in msDNA-Sen2 biosynthesis. Retron-Sen2 is an operon containing *msrmsd* (msrmsd-RNA), STM14_4640 (STM3845), and *rrtT* (STM14_4641; STM3846; RT-Sen2). *rnhA* (coding for Ribonuclease H) and *xseA*/*xseB* (coding together for Exodeoxyribonuclease VII – Exo VII) are not genetically linked to the Retron-Sen2, or to each other. **(B)** Pathway of msDNA-Sen2 production. RT-Sen2 binds to msr-RNA and reverse transcribes msd-RNA to msd-DNA, while ribonuclease H (RNase H) degrades the msd-RNA template. msr-RNA and msd-DNA are joined by a 2’-5’ phosphodiester bond (RNA-DNA 2’-5’), and by RNA-DNA hydrogen bonds at the ends of msr-RNA/msd-DNA. Exo VII cleaves the first four nucleotides off the start of the msd-DNA, separating the DNA/RNA hybrid into msDNA-Sen2 and msr-Sen2. msDNA-Sen2 remains complexed with RT-Sen2. **(C)** Strains Δ*rrtT* and Δ*xseA* are cold-sensitive. 1536 colony-arrays of the *S*Tm gene-deletion library ^22^ were pinned onto LB plates, and strains were grown at room temperature. Colony sizes ^34^ were used to calculate a fitness S-score for each strain 35. S-scores were calculated from *n* = 8. Dashed vertical line denotes the mean fitness S-score calculated from all strains (*n*=3781); negative and positive S-scores indicate sensitive and resistant mutants, respectively. **(D)** Perturbing msDNA-biogenesis leads to cold sensitivity. *S*Tm retron-deletion strains were grown for 5-6 hours at 37°C in LB, serially diluted, and spotted on LB plates. Plates were incubated either at 15 °C and 37 °C. Representative data shown from four independent experiments.

The bottleneck to understand the natural function of retrons has been the absence of phenotypes associated with retron deletions. The first retron-deletion phenotype was reported for retron-Sen2 of *Salmonella enterica* subsp. enterica ser. Typhimurium str. 14028s (*S*Tm), wherein the RT-Sen2 was found to be essential for *S*Tm survival in calves ^18^. This was because it allows *S*Tm to grow in anaerobic conditions, present in calf intestines ^19^. Here, we report that retron-Sen2 deletion mutants are also unable to grow at lower temperatures. By exploiting the retron cold-sensitivity phenotype, we show that the retron-Sen2 accessory gene STM14_4640 (*rcaT*) encodes a bona fide toxin. Perturbing msDNA biosynthesis at any stage results in toxin activation, and thereby, growth inhibition in anaerobic conditions and cold. Although reminiscent of Toxin/Antitoxin (TA) systems, which are composed of a protein or RNA antitoxin and cognate toxin ^20^, retron-Sen2 forms a novel tripartite TA system: RcaT is the toxin, and the RT-msDNA complex is the antitoxin. Using another retron encoded by *E. coli* NILS-16 ^21^, retron-Eco9, we demonstrate that this TA function is conserved and that the RT provides specificity to the TA system. We propose that bacterial retrons function as TA systems, where the RT-msDNA antitoxins directly inhibit retron-encoded toxins, by forming inactive msDNA-RT-toxin complexes.

## RESULTS

### Perturbations in msDNA-Sen2 biosynthesis inhibit *S*Tm growth in cold

As part of a larger chemical-genetics effort, we profiled the fitness of a single-gene deletion *S*Tm library ^22^ across hundreds of conditions (unpublished data). The two gene deletions that led to the highest growth sensitivity at room temperature (cold-sensitivity) were Δ*rrtT* (ΔRT-Sen2) and Δ*xseA* (Fig. 1C). Both RT-Sen2 and Exo VII are involved in msDNA-Sen2 biogenesis ^14,18^ (Fig. 1A-1B). To validate these results and exclude confounding effects (secondary mutations or polar effects), we transferred these deletions, as well as that of the accessory gene STM14_4640 (Fig. 1A), to a clean *S*Tm genetic background and flipped out the antibiotic resistance cassettes. We also constructed new marker-less deletions for the remaining msDNA-Sen2 biosynthesis genes that were not part of the library (*rnhA*, *xseB*, and *msrmsd*; Fig. 1A-B). All gene deletions, except for ΔSTM14_4640, led to severely restricted growth at 15°C (Fig. 1D) and temperatures up to 25°C (ED Fig. 1A). Consistent with previous data on Δ*rrtT* and Δ*msd* ^18^, all mutants, except for ΔSTM14_4640, also grew slower under anaerobic conditions at 37°C (ED Fig. 1B-C). It is known that msDNA-Sen2 cannot be produced in Δ*rrtT* and Δ*msrmsd* strains ^18^. We observed that RNAse H (*rnhA*) and Exo VII (*xseA*/*xseB*), but not STM14_4640, were also required for biosynthesis of mature msDNA-Sen2 (ED Fig. 1D). In summary, any perturbation in the msDNA-Sen2 biosynthesis pathway impairs *S*Tm growth at lower temperatures and anaerobiosis.

### Retron-Sen2 encodes a toxin, which is activated upon msDNA biogenesis perturbations

To understand why retron-Sen2 mutants are cold-sensitive, we isolated and sequenced spontaneous suppressors of Δ*rrtT*, Δ*xseA*, and Δ*msrmsd* mutants, that restored growth at 15°C. 28 out of 29 suppressor mutations mapped in STM14_4640, the retron-encoded accessory gene (Fig. 2A). Mutations included frameshifts (14/29), premature termination sites (4/29), or single amino acid substitutions (10/29) in STM14_4640 (ED Fig. 2A). One residue, D296, was mutated to different amino acids in all three mutant backgrounds, suggesting that it is key for the function of STM14_4640. The only suppressor that did not map in STM14_4640, had a 62 base-pair deletion within the *msd* region (Δ62:*msd*) (ED Fig. 2A). To identify the minimal *msd* deletion-region that fully suppressed cold-sensitivity, we made progressive scarless deletions in *msd* (ED Fig. 2B). Deleting up to the 63 base-pair in *msd* (Δ63:*msd*) was able to revert the cold-sensitivity phenotype (ED Fig. 2C). However, all constructs that (partially) alleviated the cold-sensitivity, resulted also in down-regulation of the expression of STM14_4640 (ED Fig. 2D), which lies directly downstream. Hence cold-sensitivity was linked in all cases to STM14_4640 integrity/expression. Therefore, we reasoned that in the absence of msDNA-Sen2, the product of STM14_4640 impairs growth at cold and anaerobic conditions. Indeed, deleting STM14_4640 in Δ*xseA*, Δ*xseB*, or Δ*rnhA* mutants, restored growth at cold temperatures (Fig. 2B) and in anaerobic conditions (Fig. 2C). We thus renamed STM14_4640 as *rcaT* (<u>r</u>etron <u>c</u>old-<u>a</u>naerobic <u>T</u>oxin).

**Figure 2.**
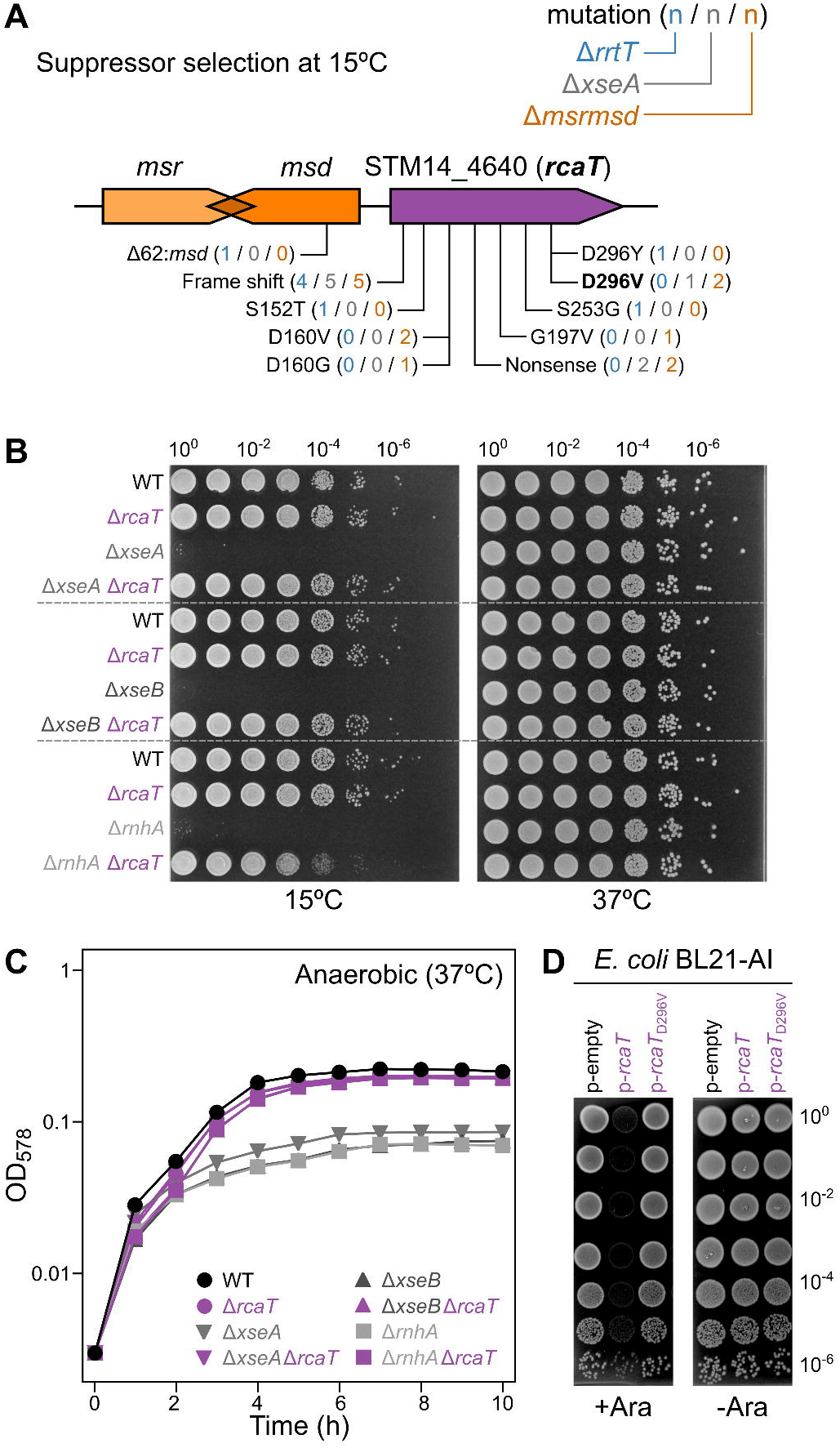
RcaT is a toxin that causes the retron phenotypes. **(A)** Cold-sensitivity suppressor mutations map almost exclusively in *rcaT*. Suppressors from *S*Tm strains (Δ*rrtT*, Δ*xseA*, Δ*msrmsd*) were isolated from LB plates incubated at 15°C, their genomic DNA was sequenced and compared to that of wildtype *S*Tm to map the mutations. **(B)** Deleting *rcaT* (STM14_4640) reverts the cold-sensitivity of retron mutants. *S*Tm strains were grown for 5-6 hours at 37°C in LB, serially diluted, and spotted on LB plates. Plates were incubated either at 15°C or 37°C. Representative data shown from two independent experiments. **(C)** Deleting *rcaT* reverts the anaerobic sensitivity of retron mutants. Growth curves of *S*Tm strains were obtained from measuring OD_578_ in microplates under anaerobic conditions at 37°C. Each point is the average optical density (OD_595_) of *n* = 11 (technical replicates), error bars denote standard deviation (error bars not shown if smaller than symbols). **(D)** Overexpressing *rcaT* is toxic in *E. coli* at 37°C. *E. coli* BL21 Arabinose-inducible (BL21-AI) carrying plasmids p-*rcaT*, or p-*rcaT*-D296V, or an empty vector (p-empty) were grown for 5-6 hours at 37°C in kanamycin-LB, serially diluted, spotted on kanamycin-LB plates with or without arabinose, and then incubated overnight at 37°C. Representative data shown from two independent experiments.

In the absence of msDNA, RcaT inhibited growth at 15°C (ED Fig. 3A), and its effect was bacteriostatic (ED Fig. 3B). Consistent with RcaT acting as a bona fide toxin, ectopically expressing RcaT, but not its inactive RcaT-D296V form, was toxic in *Escherichia coli* even at 37°C (Fig. 2D). This toxicity was exacerbated at lower temperatures (ED Fig. 3C), likely reflecting an inherent property of RcaT to be more active in cold. In these conditions, overexpressing RcaT was bactericidal (ED Fig. 3D).

Thus, growth effects that stem from impairing msDNA biosynthesis can be rescued by inactivating or down-regulating the retron-Sen2 accessory gene (*rcaT*), which encodes a toxin. This configuration is reminiscent of TA systems, where in the absence of the antitoxin, the adjacently encoded toxin inhibits bacterial growth. Consistently, toxins of TA systems are bacteriostatic at natural expression levels, but lead to killing when over-expressed ^23^.

### Retron-Sen2 is a tripartite TA system with RT and msDNA forming the antitoxin unit

Consistent with the retron-Sen2 being a novel TA system that encodes both the toxin and antitoxin activity, overexpressing the entire *S*Tm retron-Sen2 in *E. coli* (*msrmsd*-*rcaT*-*rrtT*; p-retron) did not impact growth (ED Fig. 4A), as opposed to overexpressing RcaT alone, which was toxic (Fig. 2D). RcaT toxicity was also blocked by co-expressing *msrmsd*-*rrtT in trans* from a separate plasmid (ED Fig. 4B), excluding that the antitoxin activity is dependent on *in cis* regulation. Thus, the *msrmsd* and the RT-Sen2 are sufficient to counteract the RcaT toxicity, and the entire retron constitutes a TA system.

To delineate which retron component acts as the antitoxin (branched/mature msDNA, RT, msrmsd-RNA), we constructed plasmids with retrons that lacked a single component (p-retron-Δ*rrtT*, Δ*msrmsd*, *msrmsd*^mut^, and Δ*rcaT*). The *msrmsd*^mut^ carries a single point mutation in the *msr* region, that abrogates the production of msDNA, but does not affect msrmsd-RNA expression (branching G mutation) ^11^. Additionally, we used RNAse H (Δ*rnhA*) and Exo VII (Δ*xseA*/Δ*xseB*) *E. coli* mutants to stop the msDNA biosynthesis at different steps. In all cases, deleting any retron component (except for Δ*rcaT*) or any msDNA-biosynthesis-related gene, led to the same toxicity as overexpressing *rcaT* alone (ED Fig. 4A & C). Thus, the antitoxin activity against RcaT requires the presence of all retron-components and/or the mature msDNA-Sen2.

To test whether the retron-antitoxin is counteracting RcaT by down-regulating its expression, we Flag-tagged *rcaT* in *S*Tm WT and antitoxin deletion strains, and quantified RcaT levels at 37°C and 20°C. RcaT levels remained similar in all mutants, and if anything, only slightly decreased in the Δ*rrtT* background (ED Fig. 5A-B). This is presumably due to mild polar effects of removing *rrtT*, which is located directly downstream of *rcaT*. In all cases, RcaT-3xFlag was fully functional as toxin (ED Fig. 5C). Therefore, RcaT expression is not inhibited by the retron-antitoxin, which would have led to higher RcaT levels in the mutants.

Having excluded that RcaT is counteracted by the retron-antitoxin at an expression level, we tested if this occurs by direct protein-protein interactions, as in type II TA systems. To do so, we affinity purified chromosomally encoded RT-3xFlag or RcaT-3xFlag, in wildtype and retron antitoxin-deletion *S*Tm strains (Δ*xseA*, and Δ*msrmsd*/Δ*msd*), at 37°C and 20°C, and quantified interacting proteins by quantitative mass-spectrometry (AP-qMS). Indeed, RT and RcaT strongly and reciprocally pulled down each other (Fig. 3 & ED Fig. 6). Notably, the RT-RcaT interaction occurred independently of the presence/maturation of msDNA, or of the temperature the toxin is active in. Tagged RT-3xFlag was fully functional in inhibiting RcaT (ED Fig. 7), and did not alter RT or RcaT protein levels in the input samples, compared to WT (ED Fig. 8A). In contrast, tagging RcaT led to lower RT levels in the input samples (ED Fig. 8B), but RcaT-3xFlag remained functional as toxin (ED Fig. 5C). This likely explains the lower levels of RT enrichment in the RcaT-3xFlag pull-downs (Fig. 3C). Thus, RcaT and RT stably interact with each other independently of the presence or maturation of msDNA, and of temperature.

**Figure 3.**
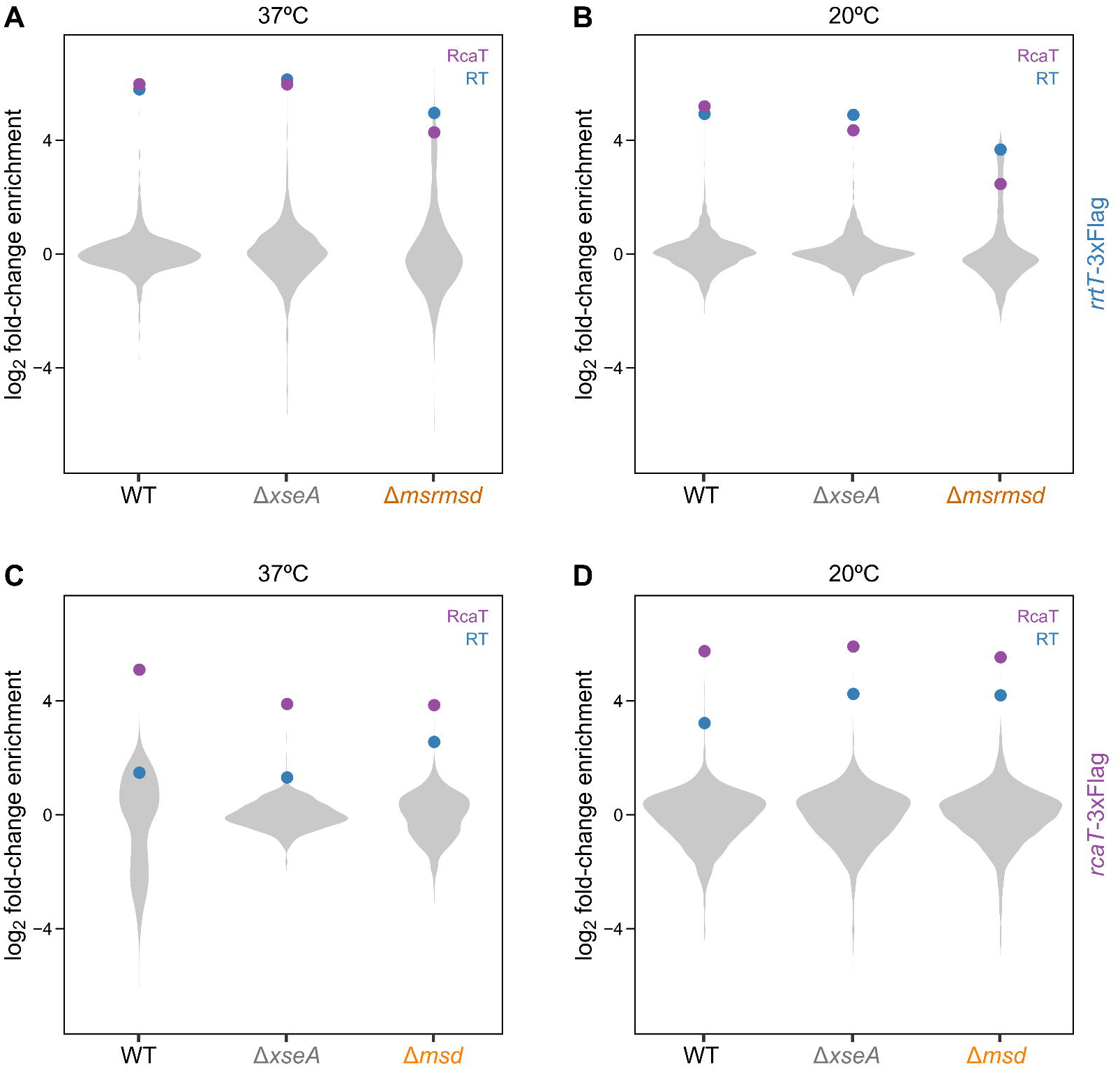
RT and RcaT reciprocally co-immunoprecipitate independent of the presence of msDNA or temperature. Immunoprecipitation of RT-3xFLAG **(A-B)** and RcaT-3xFLAG **(C-D)** at 37°C **(A, C)** and 20°C **(B, D)**. Chromosomally tagged *rrtT*-3xFlag or *rcaT*-3xFlag *S*Tm strains (WT, Δ*xseA*, Δ*msrmsd*) grown at 37°C (and for **B**, **D** shifted for 5 hours at 20°C) were used for AP-qMS. Protein abundances of IP samples of the strains carrying the tagged protein are compared to IP samples of untagged *S*Tm (y-axis). Data shown are the average from two biological replicates.

Although RcaT and RT interact in the absence of msDNA, mature msDNA production is essential for antitoxin activity. We thus wondered whether msDNA-protein interactions are involved in the antitoxin activity. Retron-RTs from different species have been previously shown to co-purify with their mature msDNA products ^15,16^. In order to assess whether RT-Sen2 also interacts with msDNA-Sen2, we first purified an RT-Sen2-6xHis protein fusion upon concomitant *msrmsd*-Sen2 expression in *E. coli* (ED Fig. 9A). The RT-Sen2-6xHis version was functional, as it could counteract the RcaT toxicity (ED Fig. 9B). At a second stage, we isolated total DNA from the purified RT-Sen2-6xHis protein sample, which yielded both mature and unprocessed msDNA-Sen2 (ED Fig. 9C). Therefore, the RT-Sen2 and msDNA-Sen2 interact with each other and are required together for the antitoxin activity.

### The RT confers specificity to the antitoxin & the msDNA modulates the antitoxin activity

Antitoxins of TA systems are specific against their cognate toxins. Since the retron-antitoxin is composed by both the RT and msDNA, we wondered which part provides the antitoxin specificity. To address this, we reasoned we could use a different retron-TA, with evolutionary diverged retron-components, and swap the individual components between retrons to make retron chimeras. For this, we used a novel retron from *E. coli* NILS-16, a clinical *E. coli* isolate ^21,24^, that we named retron-Eco9. Retron-Eco9 has an RT and an accessory gene, which are 49% and 43% identical to RT-Sen2 and RcaT-Sen2 at the protein level, respectively. *msr*-Eco9 and *msd*-Eco9 are 85% and 58% identical to their Sen2 counterparts at the nucleotide level (ED Fig. 10A), and the msDNA-Eco9 retains a similar overall structure to msDNA-Sen2 (ED Fig. 10B). As for retron-Sen2, expressing *rcaT*-Eco9 inhibited the growth of *E. coli*, while expressing the entire retron-Eco9 did not (ED Fig. 10C). In addition, the msDNA-Eco9 required RNase H/ExoVII for its production and function (ED Fig. 10D-E). Thus, the retron-Eco9 encodes a TA system similar to retron-Sen2, and its toxin (RcaT-Eco9) is inhibited by a retron-encoded antitoxin (RT-msDNA-Eco9).

To assess which part of the antitoxin unit is cognate to the toxin, we constructed chimeric retron constructs between Sen2 and Eco9. Retron-RTs are known to be highly specific in reverse transcribing their cognate msrmsd-RNA, by binding to specific RNA-structures of the msr region ^25^. Cross-specificity between non-cognate RT-msrmsd pairs has only been observed between highly homologous retrons ^13^. Thus, we first evaluated if the Sen2 and Eco9 RTs could transcribe their non-cognate msrmsd. To assess this, we isolated msDNA from *E. coli* co-expressing P_BAD_-RT plasmids (Sen2 [Se], or Eco9 [Ec]) with Ptac-msrmsd plasmids (Se, Ec, or -). Both RT-Sen2 and RT-Eco9 could use their non-cognate msrmsd and produce msDNA (Fig. 4A), albeit RT-Eco9 was slightly less efficient (lane Ec-Se – see also ED. Fig 10F for repercussions of this lower efficiency).

**Figure 4.**
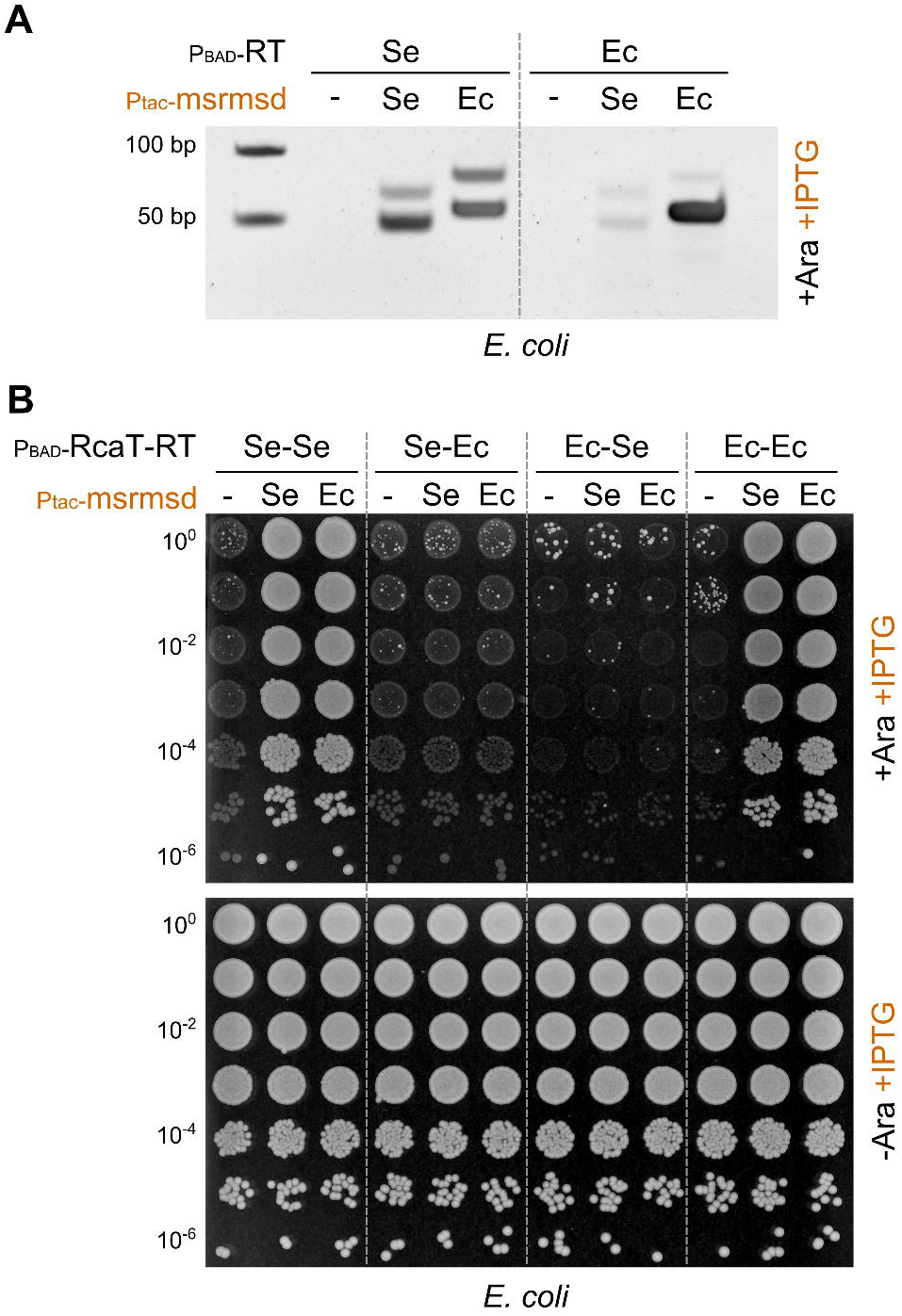
RT-RcaT interactions determine the specificity of the RT-msDNA antitoxin. **(A)** Non-cognate RT-msrmsd pairs produce msDNA. msDNA were extracted from *E. coli* BW25113 co-expressing RT of retron-Sen2 (Se) or -Eco9 (Ec) (P_BAD_-RT; Se, or Ec – arabinose induction), and plasmids carrying msrmsd (Ptac-msrmsd; Se, or Ec – IPTG induction), or an empty vector (−). Extracted msDNA were electrophoresed in a TBE-Polyacrylamide gel. Representative data shown from two independent experiments. **(B)** RcaT toxicity is inhibited by the cognate RT loaded with msDNA – cognate or not. *E. coli* BW25113 was co-transformed with plasmids carrying RT-RcaT combinations of retron-Sen2 (Se) and -Eco9 (Ec) (P_BAD_-RT-RcaT; Se-Se, Se-Ec, Ec-Se, or Ec-Ec – arabinose induction), and plasmids carrying msrmsd (Ptac-msrmsd; Se, or Ec – IPTG induction) or an empty vector (−). Strains were grown for 5-6 hours at 37°C in LB with appropriate antibiotics, serially diluted, spotted on LB plates with IPTG, antibiotics, and arabinose (+/−), and then incubated overnight at 37°C. Representative data shown from two independent experiments.

To assess the specificity of RcaT to RT, we then made arabinose-inducible plasmids, carrying binary combinations of both RcaT and RT from retrons Se, and Ec: Se-Se, Ec-Se, Se-Ec, and Ec-Ec. We co-expressed these plasmids with IPTG-inducible Ptac plasmids, expressing msrmsd from Sen2, Eco9, or the empty vector, to test which of the RT-msDNA systems retained the antitoxin-activity. As expected, both Sen2 and Eco9 RcaT inhibited the growth of *E. coli* in the absence of msDNA, irrespectively of the RT co-expressed (Fig. 4B). While cognate or non-cognate msDNA template could activate the antitoxin activity in cognate RT-RcaT combinations (Se-Se, or Ec-Ec) and neutralize the RcaT toxicity, this did not work in non-cognate RT-RcaT combinations (Se-Ec, or Ec-Se) (Fig. 4B, ED Fig 10F). This demonstrates that although RTs can produce msDNA from non-cognate msrmsd templates, and the formed RT-msDNA complex is an active antitoxin, this active antitoxin can only act against its cognate toxin. Thus, the RT-RcaT interactions (Fig. 3) are cognate within retron-TAs, and essential for antitoxin-activity.

In summary, the RT-RcaT interaction provides the specificity for the TA system, as the RT can interact directly with RcaT in the absence of msDNA, and it cannot be exchanged for a homologous RT from another retron. In contrast, the msDNA sequence is interchangeable, at least to some extent, but the RT-msDNA interaction is absolutely required to form an active antitoxin unit.

## DISCUSSION

High-throughput reverse-genetics approaches have revolutionized the characterization of gene function in bacteria ^26,27^, providing rich phenotypic information and genetic links that can be used to identify the enigmatic function of fully uncharacterized genes or to further our understanding of known cellular processes ^28–30^. Here, we have used the gene-deletion phenotypes related to the retron-Sen2 to show that retrons encode a novel family of tripartite TA systems. In retrons, the accessory gene is a toxin (RcaT) and the RT-msDNA form together an antitoxin complex (Fig. 5).

**Figure 5.**
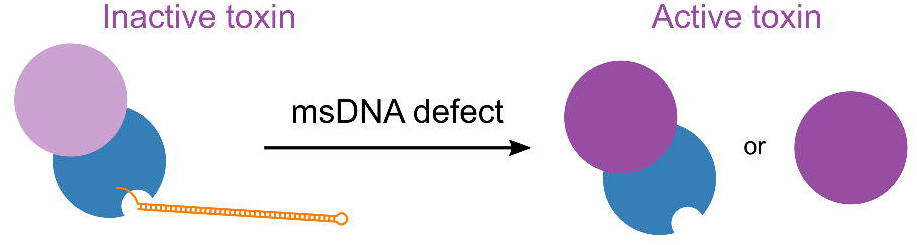
Model of retron-TA mode of action. RT-msDNA complex (antitoxin) inactivates RcaT (toxin) by direct binding. RT (blue) produces and binds msDNA (orange). The RT binds also to RcaT. Only the active antitoxin complex RT-msDNA can counterbalance the toxic activity of RcaT (light purple). RT provides the antitoxin specificity towards RcaT, but alone cannot inactive the toxin. Perturbing msDNA biosynthesis disrupts the RT-msDNA antitoxin complex, allowing for RcaT (purple) to act as toxin alone, or bound to RT.

There are a number of intriguing aspects of this new tripartite TA family that we have resolved. First, the RT is the specificity determinant of the antitoxin, by interacting with RcaT. However, alone, the RT does not affect RcaT toxicity, as the RcaT-RT complex is an active toxin (Fig. 1D, Fig. 4A), even though the RT and RcaT are tightly bound in all the conditions we tested (Fig. 3). Second, it is the processed msDNA that activates the RT-msDNA antitoxin, rather than the msrmsd-RNA or intermediate msDNA-biosynthesis forms. This finding is corroborated by the fact that when msDNA biosynthesis is stopped at steps where intermediate products become stable, then the RcaT toxin becomes active (point mutation in the branching G of *msr* stabilizing the msrmsd-RNA [ED Fig. 4A] ^11^, and *xseA*/*xseB* mutations stabilizing the unprocessed RNA-DNA hybrid [Fig. 1D, ED Fig. 4C]). Third, msDNA is required, but is not sufficient for antitoxin activity. This last point is harder to establish, as msDNA is not produced in cells lacking its cognate RT. We bypassed this limitation by using a second retron, to make hybrid-retrons. Although non-cognate RT-msrmsd pairs produce msDNA (Fig. 4A), and these form active antitoxins against cognate RT-RcaT pairs, they were not enough to inhibit the non-cognate RT-RcaT pairs (Fig. 4B). Therefore, RTs are needed for the antitoxin activity, not only because they produce msDNA, but also because they provide the antitoxin specificity by simultaneously binding the toxin (Fig. 3) and the msDNA (ED Fig. 9C).

There are also several open aspects of this retron-TA model. Although we provide strong evidence that the RT-msDNA interaction activates the antitoxin unit, the exact mechanism remains to be resolved. It is still possible that the msDNA also interacts with RcaT in the tripartite RcaT-RT-msDNA complex. In addition, we do not know the cellular target of RcaT, or why it preferentially inhibits growth in cold and anaerobic conditions, and only upon overexpression it impacts growth at 37°C in aerobic conditions. The two phenotypes may be linked, due to the target of the toxin being more relevant in cold/anaerobiosis. Alternatively, the toxin itself could be post-translationally activated in these conditions. A very different toxin in *Pseudomonas putida*, GraT, also causes cold-sensitivity upon antitoxin perturbations, and toxicity when over-expressed ^31^. GraT seems to cause cold-sensitivity by attacking ribosome biogenesis ^32^. Yet, an anaerobic-sensitivity toxin phenotype has not been reported before for GraT, or any toxin of any TA system as far as we know. Therefore, it is possible that RcaT has a distinct toxicity mechanism. Finally, we show here that another retron in *E. coli* NILS-16 encodes a similar TA system, but there are myriads of retrons, out of which only a few dozen have been experimentally verified ^4^. These carry diverse retron-components and accessory genes, and it remains to be seen if all or only a subset of them encode TA systems.

TA systems are usually bipartite, and are divided in four major types, based on how the antitoxin neutralizes the toxin ^20^. Retron-TAs are similar to type II systems, since the antitoxin (RT-msDNA) seems to inhibit RcaT through a direct protein-protein interaction. Yet, they form a distinct family of tripartite TA systems, since the msDNA is absolutely required for the antitoxin function. A number of other housekeeping genes are also required for producing the mature msDNA, increasing the complexity and the dependencies of this TA system. The evolutionary advantage of retaining such complex selfish systems is hard to fathom, as the physiological role of most TA systems remains largely unknown. In the accompanying paper ^33^, we provide an insight into this question. The exact complexity of this TA system and its msDNA component seem to allow retrons to directly sense different phage functionalities, and hence protect bacteria from phage attack.

## Supporting information

Supplementary Table 1. Proteomics data of RT-3xFlag/RcaT-3xFlag pull-downs.

Supplementary Table 2. Genotypes of bacterial strains used in this study.

Supplementary Table 3. Description of plasmids used in this study.

Supplementary Table 4. Description of construction of plasmids used in this study.

Supplementary Table 5. List of primers used in this study.

## ACKNOWLEDGEMENTS

We thank the EMBL Genomics Core Facility, and especially Anja Telzerow and Vladimir Benes for preparing the whole-genome sequencing; the EMBL Protein Expression & Purification Core Facility, and especially Jacob Scheurich and Kim Remans for purifying protein RT-Sen2; Nazgul Sakenova for help for identification of retron-Eco9, and all members of the Typas lab for discussions. This work was supported by the European Molecular Biology Laboratory and the Sofja Kovaleskaja Award of the Alexander von Humboldt Foundation. AM was supported by a fellowship from the EMBL Interdisciplinary Postdoc (EI3POD) programme under Marie Skłodowska-Curie Actions COFUND (grant number 664726). JRE was supported by the National Institutes of Health (K08AI108794). HAP is supported by NIFA (NIFA 2016-11004 & 2017-08881) and DARPA. AT is supported by an ERC consolidator grant, uCARE.

## AUTHOR CONTRIBUTIONS

MMS, JRE, HAP, & AT supervised the study. JB, JRE, HAP, & AT conceived this study. JB and AT designed the experiments, and JB, AM, SGS, and CK performed them. The *S*Tm chemical-genetics screen data is from BP & MG. MG analyzed the sequencing data and FS the AP-qMS data. JB & AM designed figures, with inputs from AT. JB & AT wrote the manuscript with input from all authors.

## DATA AVAILABILITY STATEMENT

Raw reads from whole genome sequencings are available at the European Nucleotide Archive, under project number PRJEB38324. Proteomics data from protein affinity purifications can be found in Table S1, and raw data will be uploaded in an appropriate server prior to publication. All unprocessed source images are available upon request.

## ADDITIONAL INFORMATION

Supplementary information is available for this paper. Correspondence and requests for materials should be addressed to AT.

## METHODS

### Bacterial strains, plasmids, primers, and growth conditions

Genotypes of bacterial strains, plasmid description/construction strategy, and sequences of primers used in this study are listed in Tables S2-S5, respectively. Bacteria were grown in Lysogeny Broth Lennox (LB-Lennox; Tryptone 10 g/L, Yeast Extract 5 g/L, Sodium Chloride 5 g/L). LB-Agar plates (LB plates) were prepared by adding separately autoclaved 2% molten-Agar in liquid LB. All plasmid-carrying bacterial strains were streaked-out/grown/assayed with appropriate antibiotics, in order to maintain the plasmids. Plasmids carrying P_BAD_-inserts were induced with 0.2% D-arabinose. Plasmids carrying Ptac-inserts were induced with 0.1 mM Isopropyl β- d-1-thiogalactopyranoside (IPTG). Bacterial strains with chromosomally-inserted antibiotic resistance cassettes were streaked out from stocks on antibiotic-LB plates, but grown/assayed thereafter without antibiotics. Antibiotics used were Kanamycin (30 μg/mL), Ampicillin (50 μg/mL), Spectinomycin (100 μg/mL) and Chloramphenicol (20 μg/mL). Cold-sensitive strains (*S*Tm retron-Sen2 mutants) were freshly streaked-out from glycerol stocks and kept only at 37°C before every experiment, in order to avoid suppressor mutations.

### Genetic techniques

*Salmonella enterica* subsp. *enterica* ser. Typhimurium str. 14028s (*S*Tm) chromosomal deletion strains were acquired from the *S*Tm single-gene deletion library ^22^. *S*Tm strains Δ*rnhA*, Δ*xseB*, Δ*msrmsd*, and Δ*araBAD* were constructed through λ-red recombineering ^36^, with primer-design for deletions as described in ^37^. *Escherichia coli* strains WT (BW25113), Δ*xseA*, Δ*xseB*, and Δ*rnhA* were acquired from the Keio collection ^37^. All single-gene deletion strains, newly constructed and from libraries, were re-transduced in wildtype *S*Tm or *E. coli* strains by P22 and P1 transduction, respectively. Resistance cassettes from single-gene deletions were flipped-out using the yeast-flippase expressing-plasmid, pCP20 ^38^. To make double-gene deletion strains, antibiotic resistance cassette genetic deletions were transduced in deletion strains in which the first marker was already flipped out. Δ*msd* deletion strains (ED Fig. 2B, 2C, 2D) were made in two stages. First, scar-less deletions in the *msd* region were constructed on a plasmid-vector, either by amplifying and cloning *msd* deletions from the corresponding suppressors (ED Fig. 2A), or by PCR-based directed-deletions on a plasmid carrying *msrmsd* ^39^. Second, the constructed *msd* deletion-fragments were amplified, and replaced the chromosomal-WT *msrmsd* locus by *ccdB*-recombineering ^40^ (see tables S3-S4 for details on plasmids and primers used). Plasmids were transformed in *E. coli* BW25113 strains by TSS ^41^, while *E. coli* BL21 and *S*Tm strains were transformed by electroporation ^42^.

### Growth and viability curves

For measuring growth curves anaerobically, LB was pre-reduced in anoxic conditions in an anaerobic chamber (2% H_2_, 12% CO_2_, 86% N_2_; Coy Laboratory Products) for two days before use. Flat-bottomed transparent 96-well plates containing 90 μL LB were inoculated with *S*Tm strains (grown aerobically overnight at 37°C) at OD_595_=0.01, and sealed with breathable membranes (Breathe-Easy). Plates were incubated at 37°C in the anaerobic chamber (without shaking), and OD_578_ was measured periodically (EON Biotek microplate spectrophotometer). For measuring growth under aerobic conditions at 37°C, plates were instead incubated with shaking (200 rpm), and OD_578_ was measured (Tecan Safire2 microplate spectrophotometer).

For the growth curves at 15°C, overnight cultures were inoculated in flasks of LB (OD_595_=0.01) and grown in a refrigerated incubator (Infors Multitron HT) with shaking (180 rpm). Growth was monitored by measuring OD_595_. For the viability tests samples were periodically taken, serially diluted, and plated on LB-plates. Colony Forming Units per culture volume (CFU/mL) were calculated after overnight growth at 37°C. For the viability curves in ED Fig. 3D, *E. coli* p-empty and p-*rcaT* strains were inoculated in LB (OD_595_=0.01), and cultures were incubated until OD_595_=0.4 at 37°C with shaking (180 rpm). Subsequently, cultures were transferred at 15°C with shaking (180 rpm) for 30 min. Plasmids were then induced with 0.2% arabinose, and viability was monitored by periodically plating culture-samples on ampicillin LB-plates.

### Spot growth tests

Single bacterial-colonies were inoculated in 2 mL LB, and incubated over-day aerobically at 37°C in a roller drum (6 hours, until OD_595_~5). Over-day cultures were stepwise serially-diluted eight times (ten-fold) in LB (100μL culture + 900μL LB). Using a 96-pinner (V&P Scientific, catalogue number: VP 404), ~10 μL of culture dilutions were spotted on LB-plates containing appropriate antibiotics, and 0.2% arabinose if needed (arabinose was only present in plates, not in cultures). Spots in growth tests shown in Figures 1D, 2D, ED 1A, ED 2A, ED 3C, and ED 9B were spotted manually (10 μL) with a multichannel pipette. LB-plates were incubated overnight (13-15 hours) aerobically at 37°C, in a humid incubator. For cold-sensitivity growth tests, LB-plates were incubated for 36, 48, or 72 hours, at 25°C, 20°C, or 15°C, respectively.

### msDNA isolation and running msDNA in TBE-Acrylamide gels

msDNA was isolated by alkaline lysis (reagents as described in ^43^). msDNA was over-produced by over-expressing the reverse transcriptase and the *msrmsd* region (*msrmsd*-*rrtT)*, in order to be able to purify msDNA from small culture volumes (see Tables S2-S3 for strain/plasmid combinations used for each msDNA isolation). Strains were inoculated at OD_595_=0.01 in 20 mL LB, supplemented with appropriate antibiotics and 0.2% arabinose (to over-express msDNA). Cultures were incubated for 5-6 hours at 37°C with rigorous shaking. After this, cells were put on ice and approximately 10 mL was centrifuged (4,000 rpm/15 min/4°C) – after correcting for OD_595_. Pellets were washed once with ice-cold PBS, re-suspended in alkaline solution 1, transferred into 1.5 mL Eppendorf tubes, and alkaline solutions II and III were cycled as described in ^43^. After centrifugation (14,000 rpm/20 min/4°C), supernatants were extracted twice with Phenol: Chloroform: Isoamyl-Alcohol (50:48:2, pH 8), and nucleic acids were precipitated overnight at 4°C with isopropanol. Precipitated nucleic acids were centrifuged at 14,000 rpm/60 min/4°C. Pellets were re-suspended once with 1 mL of 70% Ethanol, and centrifuged again at 14,000 rpm/60 min/4°C. Pellets (msDNA extracts) were air-dried (15 minutes), resuspended in 10 μL of distilled water containing RNase A (20 μg/mL), and incubated at 37°C for 30 minutes. Samples were subsequently kept at −80°C until further use. msDNA extracts (10 μL) were electrophoresed (70 Volts, 3.5 hours) in 1x-TBE:12%-Polyacrylamide gels (with 1x-TBE buffer), and stained with ethidium bromide. 50 bp ladder was from Promega (catalogue No. G4521).

### Affinity purifications (APs)

Two biological replicates of *S*Tm Flag-tagged strains (and appropriate negative controls, i.e., strains with the same genetic background but in which no gene was tagged) were inoculated in 100 mL LB (starting OD_595_=0.02), and grown at 37°C with constant shaking (180 rpm), until OD_595_ = 1.2-1.5. Cultures were split in half, and one flask was transferred at 20°C with constant shaking (180 rpm) for 5 hours. The remaining volume was used to prepare the 37°C samples. From this stage on, samples were kept on ice. Approximately 50 mL/OD_595_=1.5 of cultures were transferred to 50 mL tubes, with culture volumes per strain being normalized based on OD to adjust for total protein-levels across strains. Cultures were centrifuged at 5,000 rpm/10 min/4°C and the supernatant was discarded. Pellets were washed once with 50 mL of ice-cold PBS, and cells were centrifuged again. Pellets were then frozen at −80°C. Subsequently, pellets were re-suspended in 1.2 mL of Lysis Buffer (50 μg/ml lysozyme, 0.8% NP-40, 1 mM MgCl_2_, 1x protease inhibitors [Roche; cOmplete Protease Inhibitor Cocktail] in PBS), and transferred to Eppendorf tubes. Cells were lysed by ten freeze-thawing cycles (frozen in liquid nitrogen, thawing 5 minutes at 25°C with 1,400 rpm shaking per cycle). Lysates were centrifuged at 14,000 rpm/60 min/4°C to remove intact cells and other insoluble components. Samples were taken at this step (input samples). Flag-beads (ANTI-FLAG^®^ M2 Affinity Agarose Gel; Sigma-Aldrich) were washed twice (20x the beads volume) with Wash Buffer (0.8% NP-40 in PBS), and 25 μL of washed Flag-beads were added to ~1 mL of lysate. Lysates were incubated with Flag-beads overnight on a table-top roller, at 4°C. Subsequently, beads were centrifuged at 8,200 rcf/10 min/RT, and the supernatants were discarded. The beads were washed four times with 1 mL of Wash Buffer (2 min rolling with table-top roller and centrifuged at 8,200 rcf/2 min/ RT per wash cycle). After the final wash, 50 μL of Elution Buffer (150 μg/mL 3xFlag peptide [Sigma-Aldrich], 0.05% Rapigest [Waters], 1x protease inhibitors in PBS) was added to the Flag-beads, and proteins were eluted for 2 hours on a table-top roller, at 4°C. The samples were centrifuged at 8,200 rcf/15 min/RT, 50 μL of eluates were retrieved, and transferred to Eppendorf tubes (IP samples).

### Proteomics analysis of APs

Proteins were digested according to a modified SP3 protocol ^44^. Briefly, approximately 2 μg of protein was diluted to a total volume of 20 μL of water and added to the bead suspension (10 μg of beads (Thermo Fischer Scientific—Sera-Mag Speed Beads, CAT# 4515-2105-050250, 6515-2105-050250) in 10 μL 15% formic acid and 30 μL ethanol). After a 15 min incubation at room temperature with shaking, beads were washed four times with 70% ethanol. Next, proteins were digested overnight by adding 40 μL of digest solution (5 mM chloroacetamide, 1.25 mM TCEP, 200 ng trypsin, and 200 ng LysC in 100 mM HEPES pH 8). Peptides were eluted from the beads, dried under vacuum, reconstituted in 10 μL of water, and labelled for 30 min at room temperature with 17 μg of TMT10plex (Thermo Fisher Scientific) dissolved in 4 μL of acetonitrile. The reaction was quenched with 4 μL of 5% hydroxylamine, and experiments belonging to the same mass spectrometry run were combined. Samples were desalted with solid-phase extraction on a Waters OASIS HLB μElution Plate (30 μm) and fractionated under high pH conditions prior to analysis with liquid chromatography coupled to tandem mass spectrometry (Q Exactive Plus; Thermo Fisher Scientific), as previously described ^45^. Mass spectrometry raw files were processed with isobarQuant, and peptide and protein identification was performed with Mascot 2.5.1 (Matrix Science) against the *S*Tm UniProt FASTA (Proteome ID: UP000001014), modified to include known contaminants and the reversed protein sequences (search parameters: trypsin; missed cleavages 3; peptide tolerance 10 ppm; MS/MS tolerance 0.02 Da; fixed modifications were carbamidomethyl on cysteines and TMT10plex on lysine; variable modifications included acetylation on protein N-terminus, oxidation of methionine, and TMT10plex on peptide N-termini).

The fold-enrichment of pulled-down proteins in Flag-tagged strains compared to negative controls was calculated, and statistical significance was evaluated using limma analysis ^46^. A similar analysis was conducted on the input samples to ensure that enriched proteins were not overexpressed in Flag-tagged strains.

### Whole genome sequencing

Genomic DNAs from retron-mutant suppressor strains were isolated using a kit, by following the guidelines of the manufacturer (NucleoSpin Tissue, Mini kit for DNA from cells and tissue; REF 740952.50). For genomic DNA library preparation, 1 μg of input DNA was fragmented with sonication for 2 minutes, and libraries were constructed using a kit (NEB Ultra DNA library kit for Illumina; catalogue number E7370L), according to the manufacturer’s instructions. The 30 genomic libraries were sequenced using a NextSeq Illumina platform with a 150 base pairs paired-end configuration. Variants were called using breseq v0.28.0 ^47^, using the *Salmonella enterica subsp. enterica* serovar Typhimurium strain 14028S genome as reference (RefSeq ID: NC_016856.1). Genotypes of suppressor strains can be found in Table S2.

### RT-Sen2 purification and DNA-isolation from RT-Sen2 purified protein

Plasmid pJB120 (pET28α-*msrmsd*-*rrtT*-His; described in Tables S3-S4) was used to over-express a C-terminally His-tagged RT-Sen2 protein, along with msrmsd-RNA. Strain *E. coli* BL21 (DE3) CodonPlus-RIL was electroporated with pJB120. An overnight culture of the overexpression strain was inoculated at OD_595_=0.01 in 100 mL of Auto-Induction Medium (ZYM-5052 ^48^), incubated at 37°C with shaking (180 rpm) until OD_595_ ~ 0.6, and then incubated at 20°C with shaking (180 rpm) overnight. The saturated culture was centrifuged (4,000 rpm/10 min/4°C), and resuspended in 40 mL of Lysis Buffer (50 mM Tris pH = 8, 500 mM NaCl, 20 mM Imidazole, 10% Glycerol). Cells were lysed by passaging them six times using a microfluidizer. Lysates were centrifuged (35,000 rpm/25 min/4°C), and cleared lysates were added to washed Nickel-beads. After discarding the flow-through, beads were washed thrice with 5 mL of Lysis Buffer, and bound proteins were eluted with 1 mL of Elution Buffer in five fractions (50 mM Tris pH = 8, 500 mM NaCl, 300 mM Imidazole, 10% Glycerol). Fractions were pooled together. 500 μg of RT-Sen2 protein were used to isolate DNA (shown in ED Fig. 9B). DNA was extracted from the elution fraction twice with Phenol: Chloroform: Isoamyl-Alcohol (25:24:1, pH 8), and then nucleic acids were precipitated overnight at 4°C with isopropanol. Precipitated nucleic acids were centrifuged at 14,000 rpm/60 min/4°C. Pellets were re-suspended once with 1 mL of 70% Ethanol, and centrifuged again at 14,000 rpm/60 min/4°C. Resultant pellets were air-dried (15 minutes), resuspended in 10 μL of distilled water containing RNase A (20 μg/mL), and incubated at 37°C for 30 minutes. The sample was loaded on a 1x-TBE:12%-Polyacrylamide gel (with 1x-TBE buffer), electrophoresed at 70V for 3.5 hours (constant voltage), and DNA was stained with ethidium bromide.

### SDS-PAGE and Immunoblot

Samples from *rrtT*-3xFlag, *rcaT*-3xFlag, and control strains were suspended in 1x Laemmli buffer, and heated to 95°C for 10 min. Proteins were separated by SDS-PAGE, and the gel was blotted to a PVDF membrane. Membranes were blocked for 1 hour (RT) with 5% skimmed milk in TBS-T (TBS-TM), and probed over-night at 4°C either in TBS-TM with a 1:1000 anti-Flag antibody (Sigma-Aldrich; catalogue No F3165), or with a 1:10000 anti-LpoA antibody (loading control ^28^; wherever applicable, membranes were cut, with one half probed with anti-LpoA, and the other half with anti-Flag). Membranes were incubated for 1 hour with HRP-conjugated secondary antibodies (1:5000, anti-mouse, Sigma-Aldrich Catalogue No A9044; Flag, or 1:10000, anti-rabbit, Merck, Catalogue No GENA934; LpoA) in TBS-TM. After washing with TBS-T, chemiluminescence substrate (GE-Healthcare) was added, and signal was detected using X-ray films (Advantsta). X-ray films were then scanned at 300×300 dpi. Digital images were cropped, and adjusted in Inkscape. Signal quantifications were done in ImageJ.

### Retron-Eco9 identification

Retron-Sen2 components (*rcaT* and *rrtT* gene products) were used as queries in pBLAST to identify retron elements in the *E. coli* natural isolate collection ^24^. Retron-Eco9 was selected as having proteins homologous to both *rcaT*-Sen2 and *rrtT*.

## EXTENDED DATA FIGURE LEGENDS

**Extended Data Figure 1.**
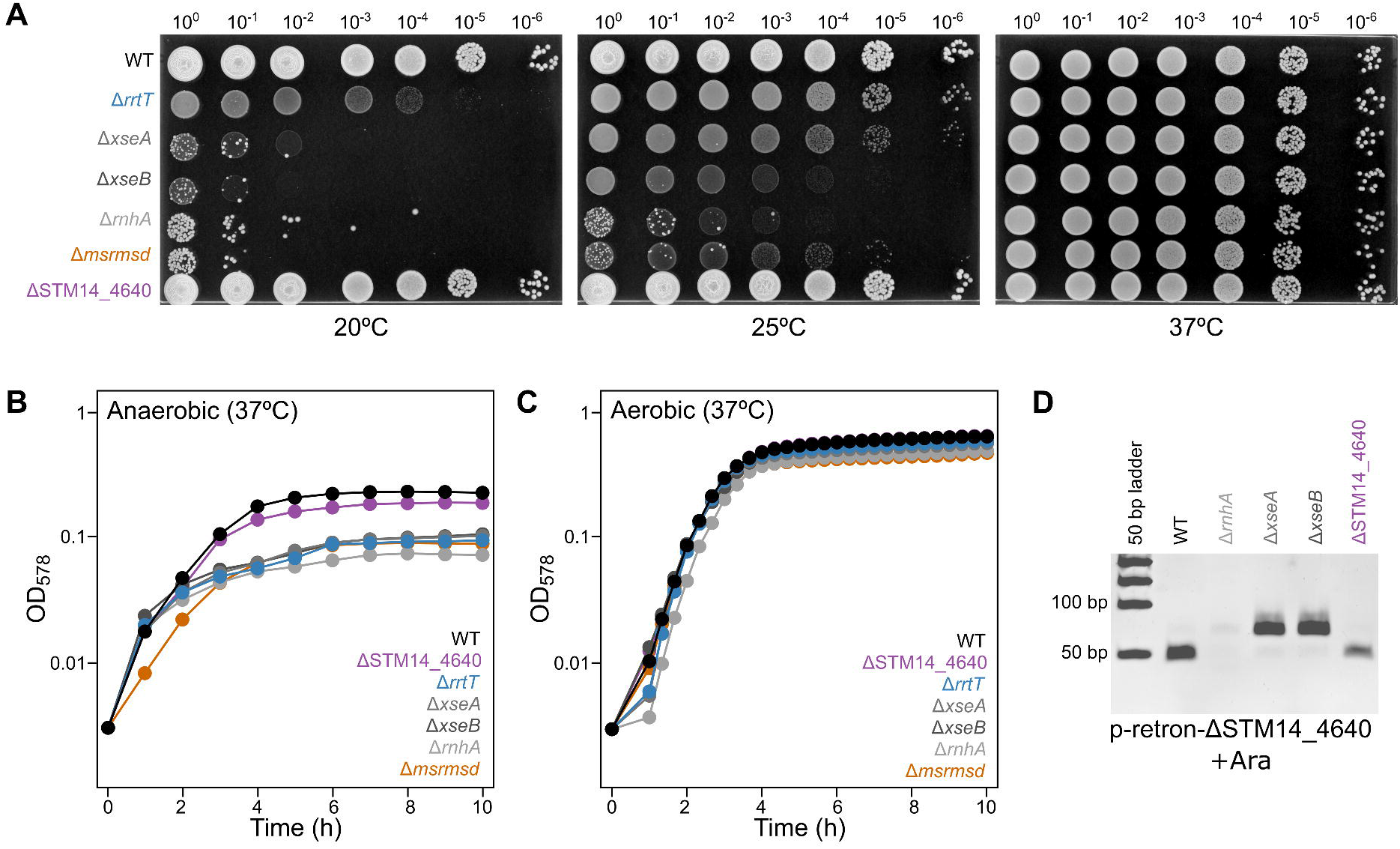
Retron phenotypes and msDNA-Sen2 biosynthesis. **(A)** Perturbing msDNA-biogenesis leads to cold-sensitivity. *S*Tm wildtype and retron-deletion strains were serially diluted and spotted on LB plates as in Fig. 1D. Plates were incubated at 20°C, 25°C, or 37 °C. Representative data shown from four independent experiments. **(B)** Retron mutants grow slower in anaerobic conditions. Growth curves of *S*Tm wildtype and retron-deletion strains were obtained by measuring OD_578_ in microtiter plates, under anaerobic conditions at 37°C. Data plotted as in Fig. 2C. **(C)** Retron mutants are not affected in aerobic conditions. Experiment as in panel B, but strains were grown aerobically. **(D)** RNAse H and Exonuclease VII are involved in msDNA biosynthesis. msDNA was extracted from *S*Tm wildtype and retron-deletion strains carrying plasmid p-retron-ΔSTM14_4640. Extracted msDNA was electrophoresed in TBE-Polyacrylamide gels. A representative gel from three independent experiments is shown.

**Extended Data Figure 2.**
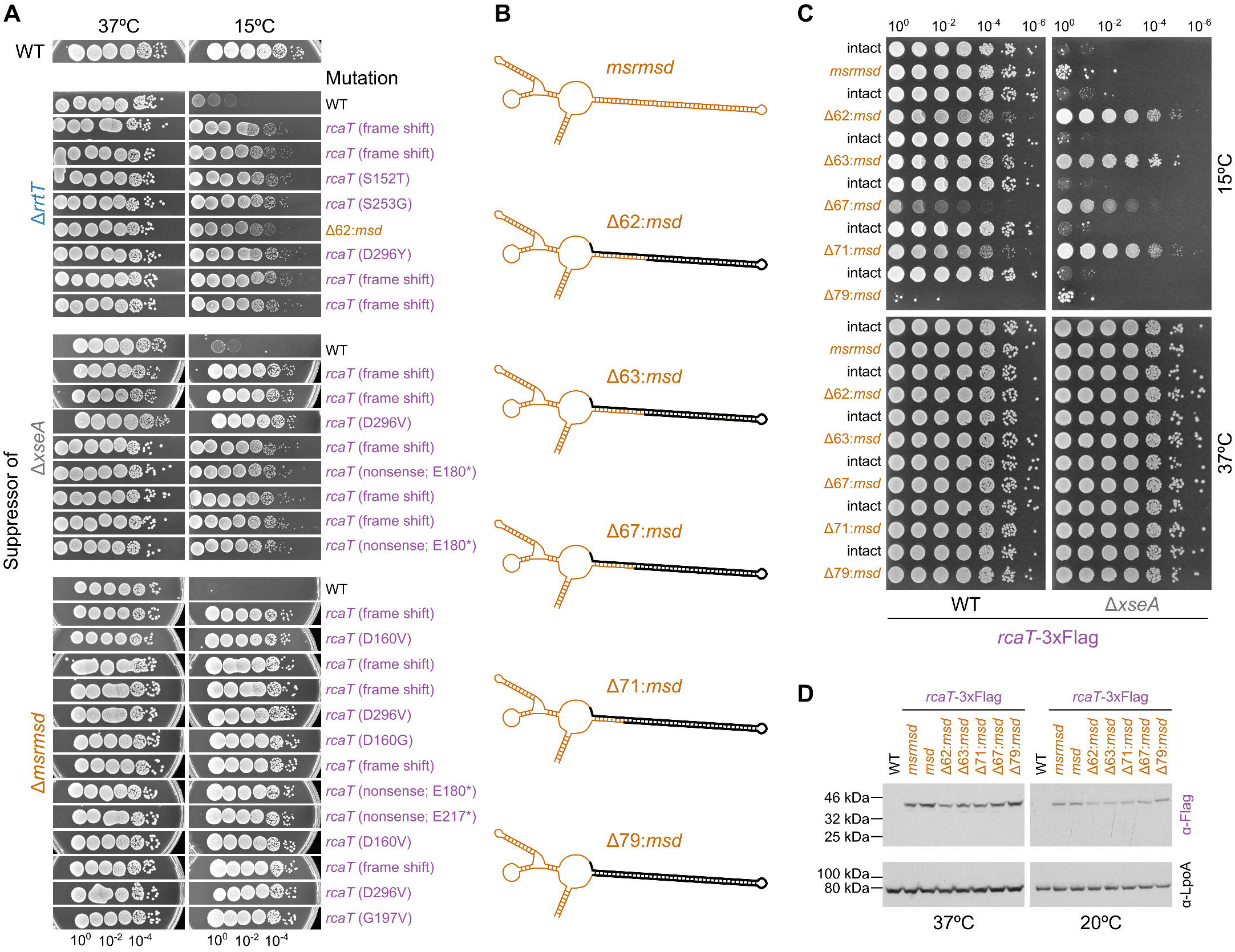
Retron phenotype suppressing mutations inactivate *rcaT*. **(A)** Suppressor strains are reverted to wildtype growth in cold temperatures. Suppressors isolated from cold-sensitive *S*Tm mutants (Δ*rrtT*, Δ*xseA*, Δ*msrmsd*) were grown, serially diluted and spotted on LB plates as in Fig. 1D. Identified suppressor mutations are indicated. **(B)** Progressive *msd* deletions constructed – in black the region deleted from the msrmsd-RNA. **(C)** Internal *msd* deletions from Δ62 to Δ71 suppress the cold-sensitivity of Δ*xseA* cells, but induce cold-sensitivity in wildtype, presumably because they are antitoxin-deficient. Wildtype (WT) or Δ*xseA S*Tm strains carrying *msd* deletions and the intact *msrmsd* region were grown, serially diluted and spotted on LB plates as in Fig. 1D. In contrast to all other internal deletions, the Δ79:*msd* mutant behaves as a full *msrmsd* deletion. Representative data shown from two independent experiments. **(D)** Internal *msd* deletions (from Δ62 to Δ71) result in lower RcaT levels. *rcaT*-3xFlag tagged *S*Tm strains carrying *msd* deletions and the WT *S*Tm strain were either grown in LB only at 37°C, or shifted to 20°C for 5 hours. Protein samples from strains were analysed by SDS-PAGE and immunoblotting. LpoA levels (α-LpoA antibody) were used as a loading control. Representative data shown from two independent experiments.

**Extended Data Figure 3.**
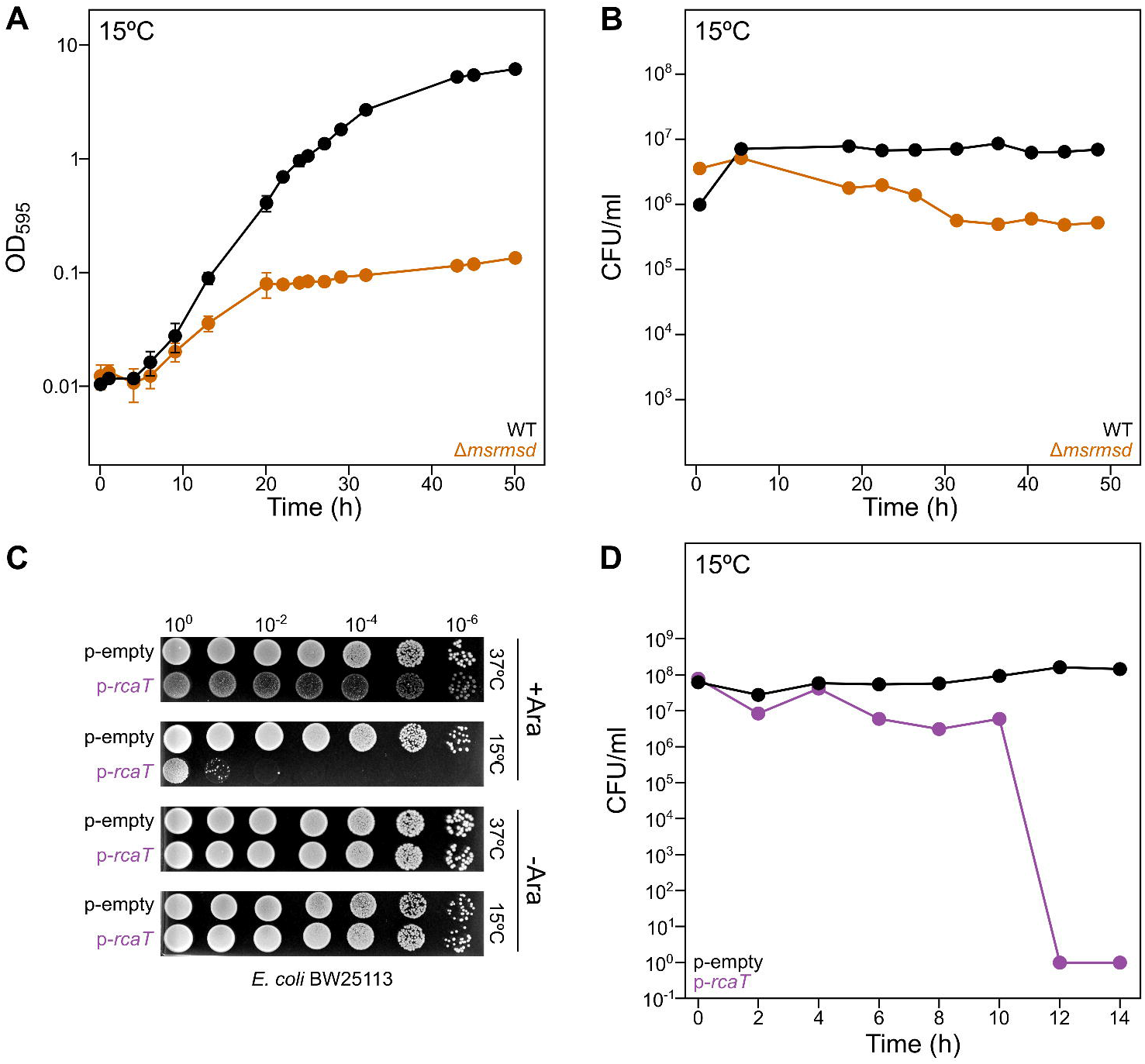
RcaT is bacteriostatic at native levels, but becomes bactericidal at higher levels. **(A)** RcaT inhibits growth at 15°C in *S*Tm. *S*Tm strains (WT and Δ*msrmsd*) were grown overnight at 37°C and used to inoculate new cultures at 15°C for which the growth as monitored by measuring optical density (OD_595_). Data points represent the average of three measurements (biological replicates). Error bars denote standard deviation (if not shown, smaller than the symbols). **(B)** RcaT is bacteriostatic at native levels in *S*Tm. *S*Tm strains (WT and Δ*msrmsd*) were grown at 15°C, and viability curves were obtained by plating culture sample dilutions on LB plates to count colony forming units (CFU) per ml. Data points represent one experiment. **(C)** Cold aggravates RcaT-mediated toxicity in *E. coli*. *E. coli* carrying plasmid p-*rcaT* or an empty vector were grown in ampicillin-LB for 5-6 hours at 37°C, serially diluted and spotted on ampicillin-LB plates with or without arabinose (Ara). Representative data shown from three independent experiments. **(D)** RcaT is bactericidal when overexpressed in *E. coli. E. coli* BW25113 strains carrying plasmid p-*rcaT* or an empty vector were grown in ampicillin-LB at 37°C, and then cultures were transferred at 15°C, and induced with arabinose. Viability curves were obtained by monitoring growth over time and plating culture samples on ampicillin-LB plates to count colony forming units (CFU) per ml. Data points represent the average of two experiments (biological replicates). Error bars denote standard deviation (if not shown, smaller than the symbol).

**Extended Data Figure 4.**
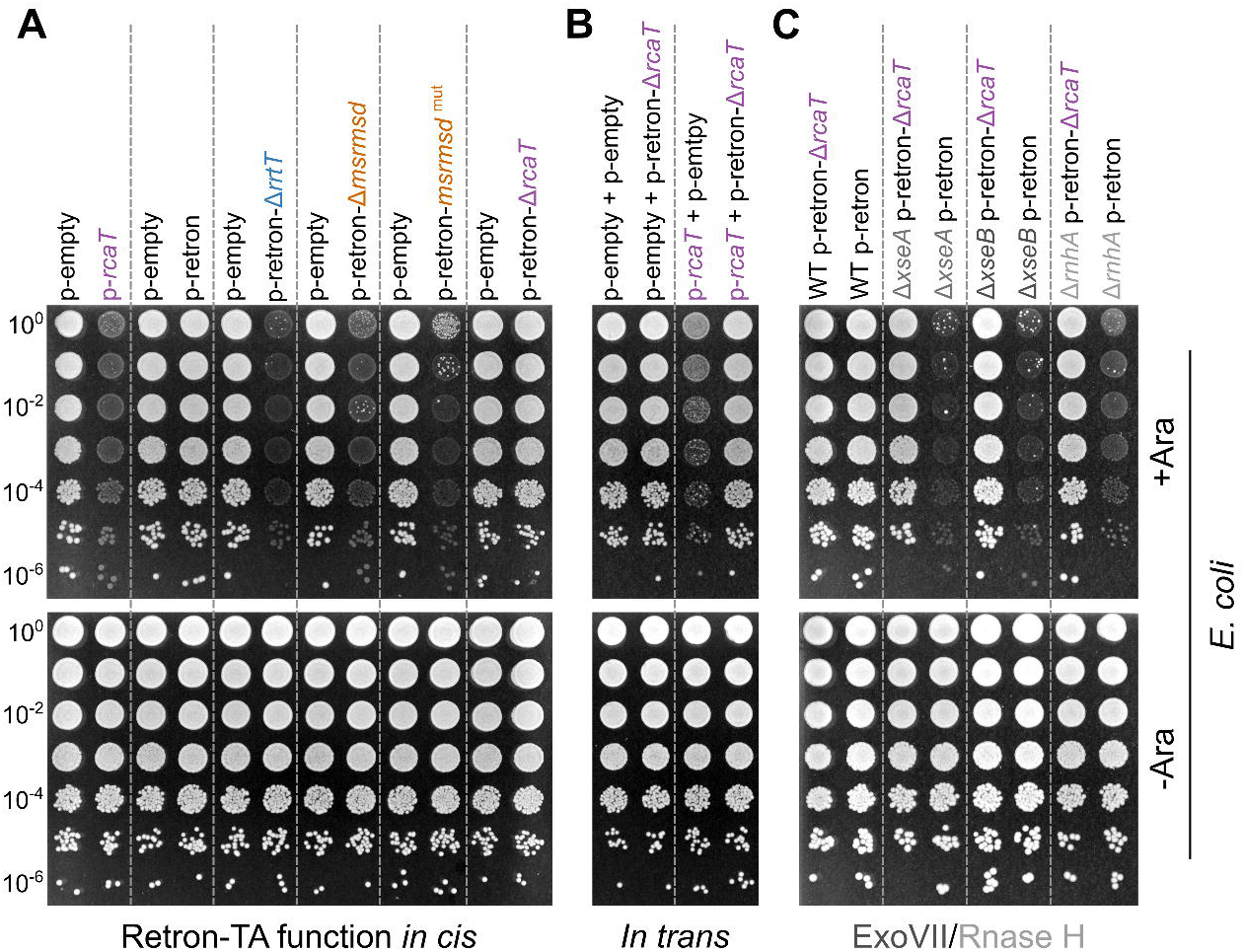
Retron functions as a toxin/antitoxin system in *E. coli*. **(A)** RcaT inhibition requires both *msrmsd* and *rrtT* in *E. coli*. *E. coli* with plasmids carrying retron-components were grown for 5-6 hours at 37°C in spectinomycin-LB, serially diluted, spotted on spectinomycin-LB plates with or without arabinose, and plates were incubated at 37°C. Only the intact retron could restore growth. RcaT expression is sufficient to inhibit growth. Representative data shown from three independent experiments. **(B)** RcaT can be inhibited by *msrmsd*-*rrtT* also *in trans*. *E. coli* carrying binary combinations of plasmids p-*rcaT*,p-retron-Δ*rcaT*, and empty vectors, were grown in LB with appropriate antibiotics, serially diluted and spotted as in **A**. Representative data shown from three independent experiments. **(C)** RcaT inhibition requires RNAse H and Exo VII in *E. coli*. *E. coli* strains (WT, Δ*xseA*, Δ*xseB*, Δ*rnhA*) carrying plasmid p-retron-Δ*rcaT* or p-retron were grown, serially diluted and spotted as in **A**. Representative data shown from two independent experiments.

**Extended Data Figure 5.**
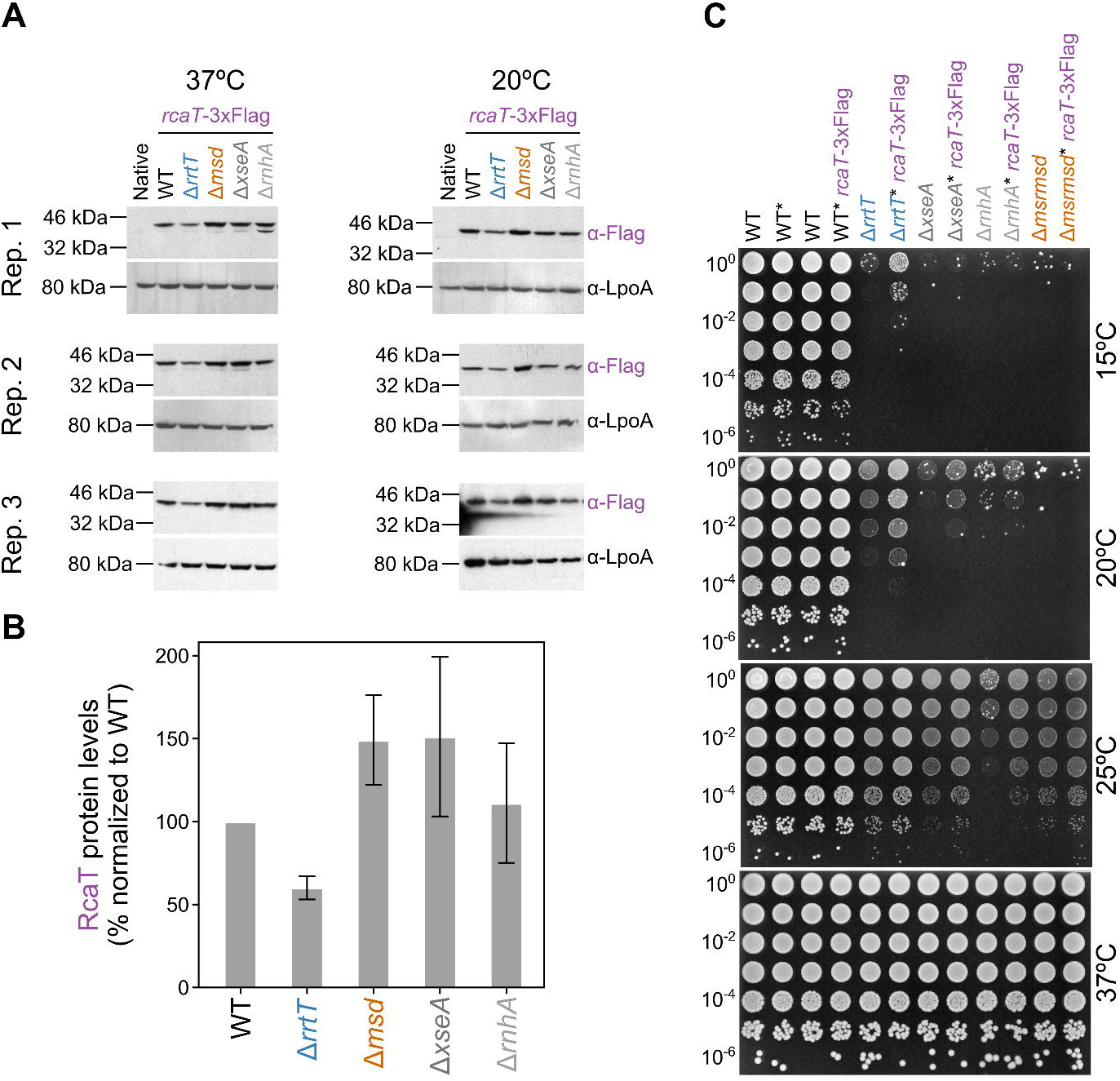
Antitoxin does not act on RcaT expression. **(A)** RcaT levels remain largely unaffected by antitoxin deletions. *rcaT*-3xFlag *S*Tm strains (WT and retron-deletions) and the *S*Tm untagged strain (native) were either grown in LB only at 37°C, or shifted to 20°C for 5 hours. Protein samples from strains were analysed by SDS-PAGE and immunoblotting. LpoA levels (α-LpoA antibody) were used as a loading control. Data shown from three independent experiments. **(B)** Quantification of RcaT-3xFlag signal from immunoblots in **A** using ImageJ (pixel-density). Error bars depict standard deviation. **(C)** Flag-Tagged RcaT retains its function. *rcaT*-3xFlag *S*Tm strains (wildtype–WT and retron-deletions) and their untagged counterparts were grown for 5-6 hours at 37°C in LB, serially diluted, spotted on LB plates, and incubated at 37°C, 25°C, 20°C, or 15°C. * denotes ΔSTM14_4645::cat, which is used to co-transduce the scarless Flag-tagged *rcaT* (co-transduction verified by PCR). Representative data shown from two independent experiments.

**Extended Data Figure 6.**
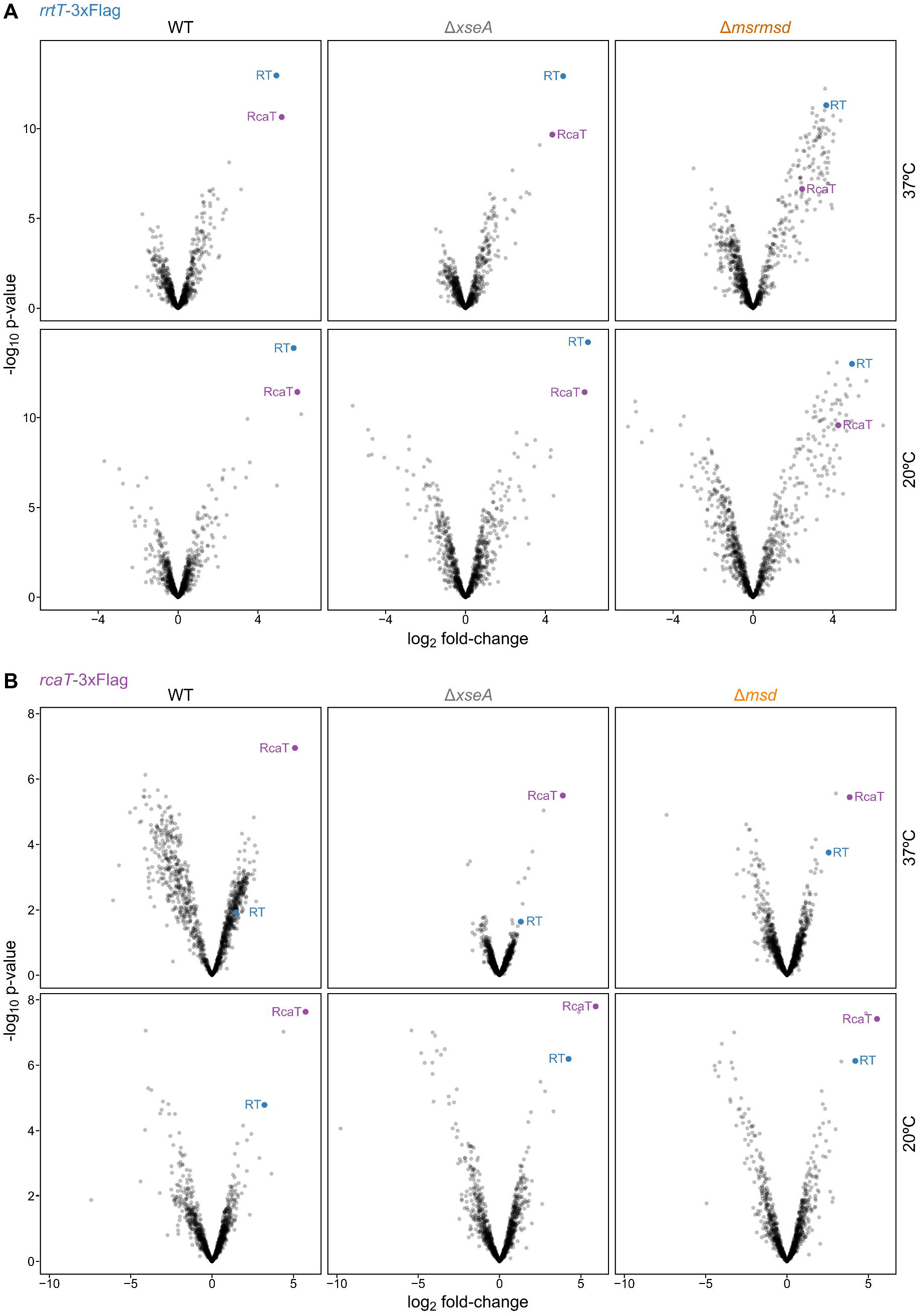
RT and RcaT reciprocally pull down each other. Volcano plots of APs of RT-Sen2 **(A)** and RcaT **(B)** at 37°C and 20°C in wildtype and different mutant backgrounds, performed as described in Fig. 3. The y-axis represents p-values of identified peptides from two biological replicates in *rrtT*-3xFlag and *rcaT*-3xFlag IP samples, and the x-axis represents the peptide-enrichment in the two samples compared to an untagged *S*Tm strain control.

**Extended Data Figure 7.**
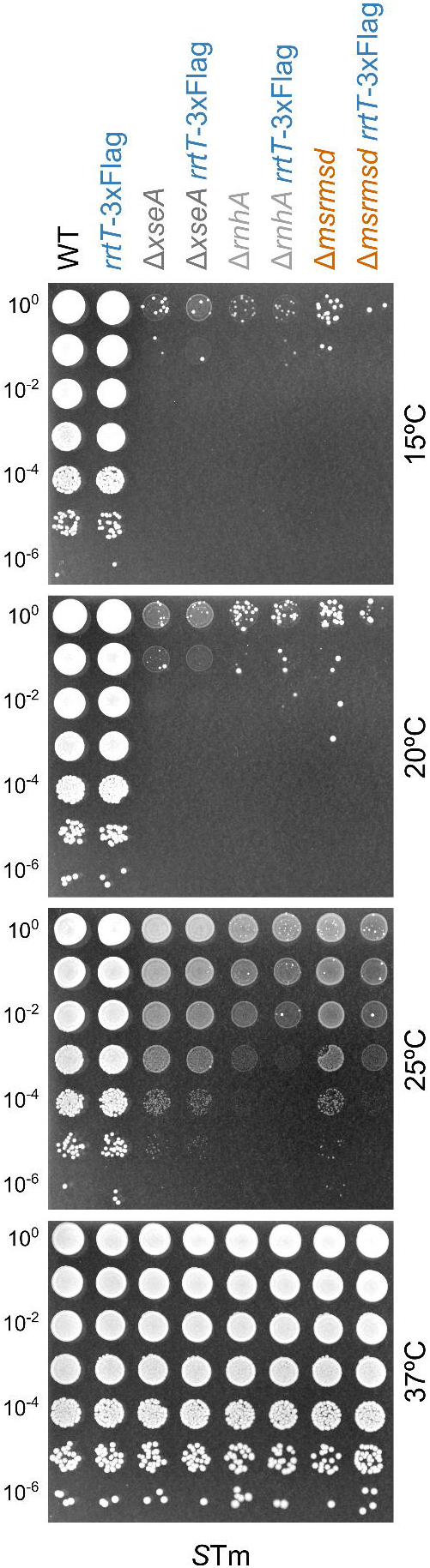
Flag-tagged RrtT retains its function. *rrtT*-3xFlag *S*Tm strains (WT and retron-deletions) and their untagged counterparts were grown for 5-6 hours at 37°C in LB, serially diluted, spotted on LB plates, and plates were incubated either at 37°C, 25°C, 20°C, or 15°C.

**Extended Data Figure 8.**
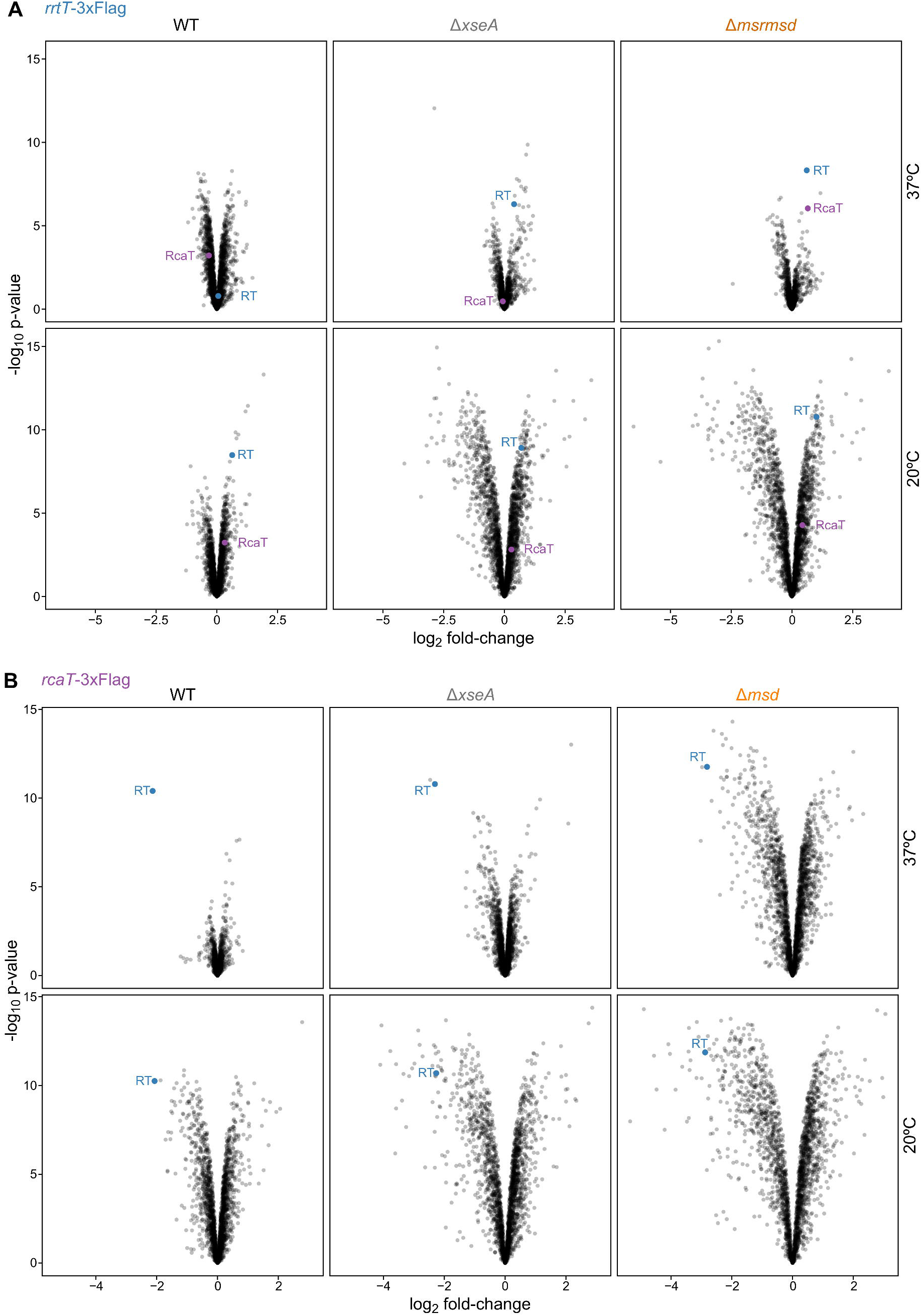
Effects of tagging on retron-protein levels. Flag-tagging *rrtT* does not alter retron protein expression **(A)**, whereas flag-tagging *rcaT* decreases retron protein levels **(B)**. Proteins in input (whole proteome) samples used for AP samples shown in Fig. 3 were quantified by mass spectrometry. Protein abundances in input samples of Flag-tagged strains were compared to input samples of untagged *S*Tm strains (x-axis). The y-axis represents the p-values of log2 fold-changes of quantified proteins. Data derived from two biological replicates.

**Extended Data Figure 9.**
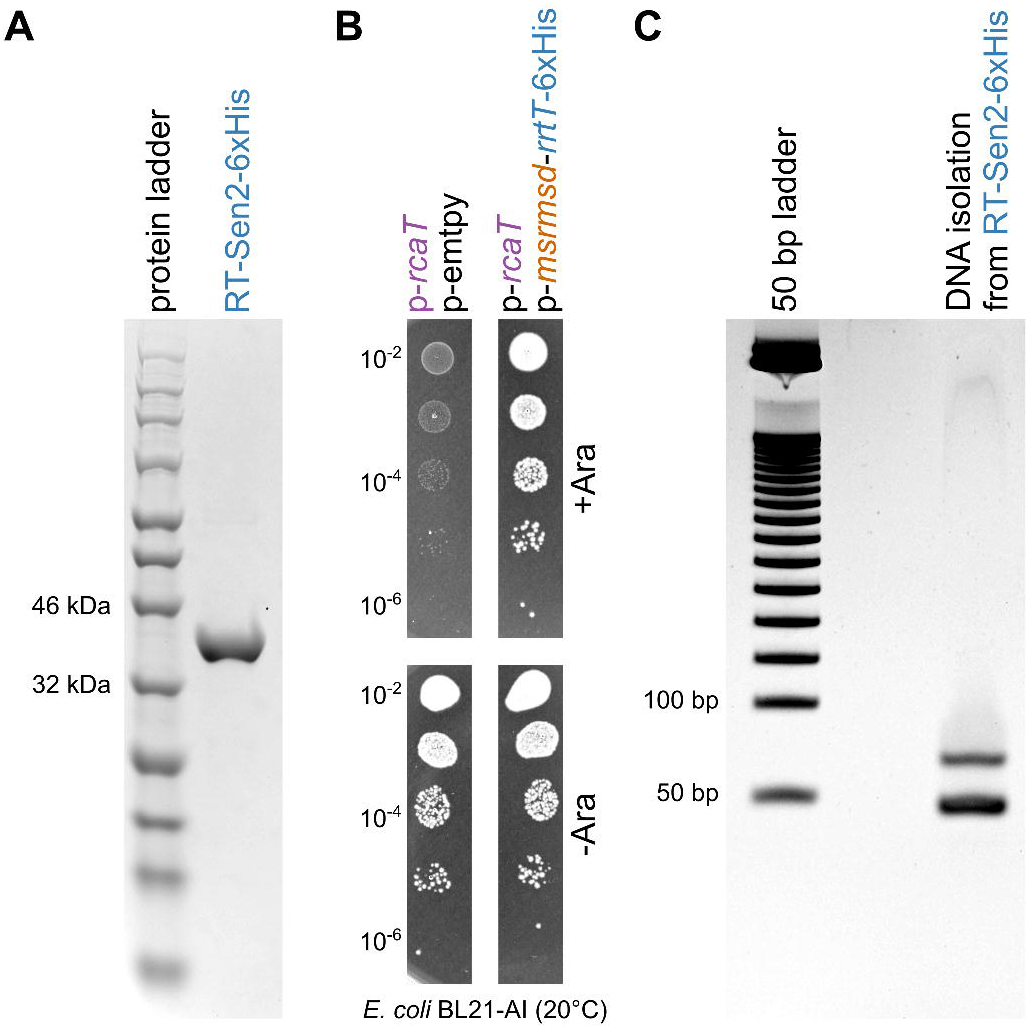
RT interacts with msDNA. **(A)** Purification of protein RT-Sen2. An *E. coli* BL21 (DE3) CodonPlus-RIL strain carrying plasmid p-*msrmsd*-*rrtT*-6xHis was used to purify protein RT-Sen2-6xHis (C-terminal fusion), by nickel-column immobilized metal-affinity chromatography. **(B)** His-tagging *rrtT* does not affect its antitoxin activity. *E. coli* BL21-AI strains carrying binary combinations of plasmids p-*msrmsd*-*rrtT*-6xHis, p-*rcaT*, and empty vectors, were grown for 5-6 hours at 37°C in kanamycin-LB, serially diluted, spotted on kanamycin-LB plates with or without arabinose, and plates were incubated at 20°C. **(C)** Isolation of msDNA from purified RT-Sen2. Total DNA was extracted from 500 μg of purified RT-Sen2-6xHis protein. Resultant DNA was electrophoresed on a TBE-Polyacrylamide gel.

**Extended Data Figure 10.**
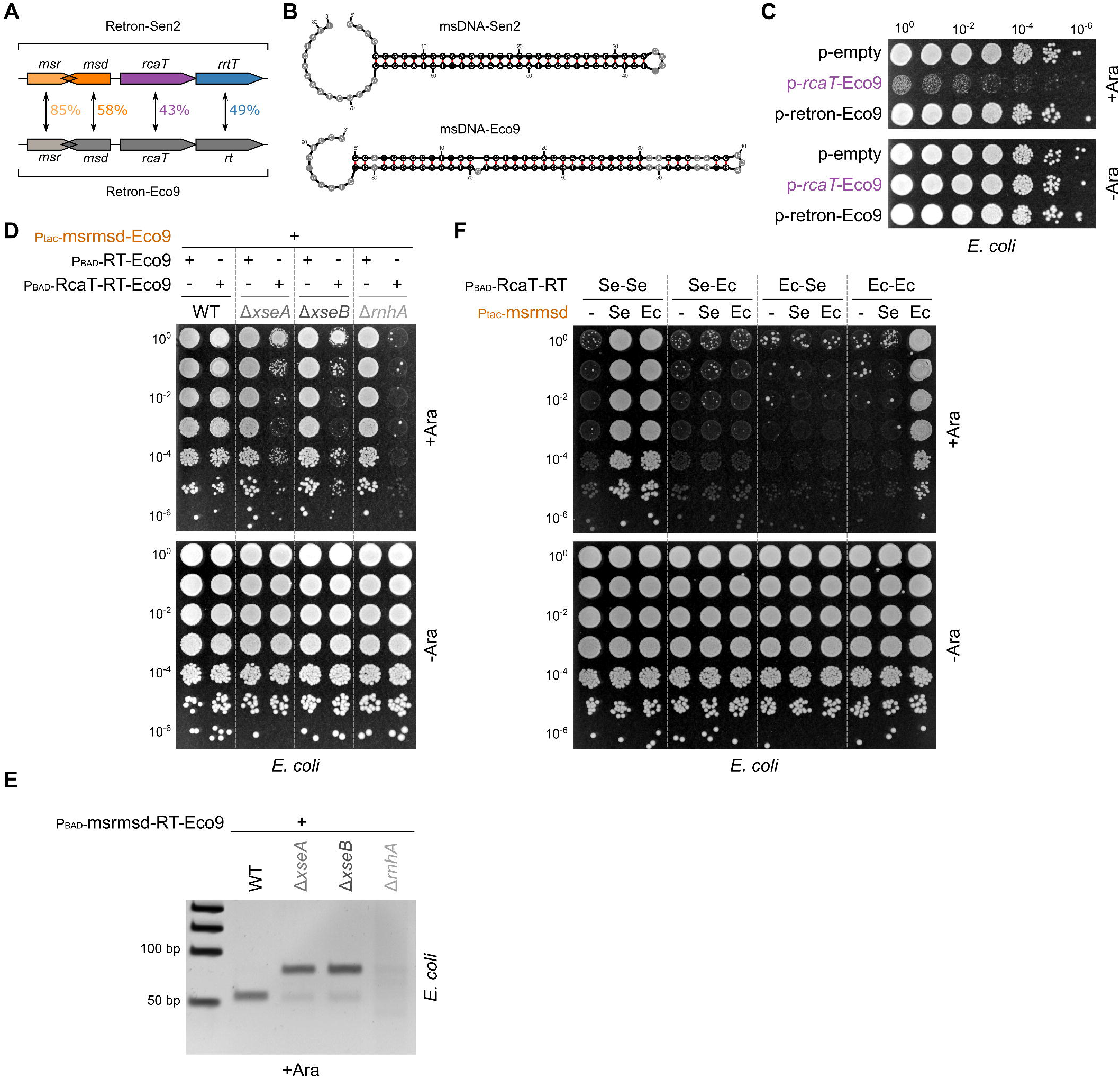
*E. coli* Retron-Eco9 is similar to *S*Tm retron-Sen2. **(A)** Retron-Eco9 has a similar operon structure to retron-Sen2. Retron-Eco9 contains *msr*, *msd*, *rcaT* and *rrtT* regions, which are 85%, 58%, 43%, and 49% identical (first two nucleotide level, last two protein level) to the corresponding retron-Sen2 regions. **(B)** msDNA-Eco9 is of similar structure to msDNA-Sen2. Models of msDNA-Sen2 and msDNA-Eco9 are depicted, created using ^49^. **(C)** Retron-Eco9 is a TA system. *E. coli* BW25113 strains carrying plasmids p-*rcaT*-Eco9, p-retron-Eco9, or an empty vector, were grown for 5-6 hours at 37°C in chloramphenicol-LB, serially diluted, spotted on chloramphenicol-LB plates with or without arabinose, and plates were incubated at 37°C. Representative data shown from three independent experiments. **(D)** RcaT-Eco9 inhibition requires RNAse H and Exo VII in *E. coli*. *E. coli* BW25113 strains (wildtype, Δ*xseA*, Δ*xseB*, Δ*rnhA*) carrying plasmid Ptac-msrmsd-Eco9, along with either P_BAD_-RT-Eco9, or P_BAD_-RT-RcaT-Eco9, were grown for 5-6 hours at 37°C in LB with appropriate antibiotics, serially diluted, spotted on LB plates with antibiotics, with/without arabinose, and plates were incubated overnight at 37°C. Representative data shown from two independent experiments. **(E)** msDNA-Eco9 biosynthesis requires RNase H and Exo VII. msDNA was extracted from *E. coli* BW25113 strains (wildtype, Δ*xseA*, Δ*xseB*, Δ*rnhA*) carrying plasmid P_BAD_-msrmsd-RT-Eco9. Extracted msDNA were electrophoresed in a TBE-Polyacrylamide gel. **(F)** The Ec RT cannot be activated as antitoxin by basal non-cognate msDNA (Se) levels (compare with Fig. 4B). *E. coli* BW25113 was co-transformed with plasmids carrying RT-RcaT combinations of retron-Sen2 (Se) and -Eco9 (Ec) (P_BAD_-RT-RcaT; Se-Se, Se-Ec, Ec-Se, or Ec-Ec – arabinose induction), and either plasmids carrying msrmsd (Ptac-msrmsd; Se, or Ec – IPTG induction) or an empty vector (−). Strains were grown for 5-6 hours at 37°C in LB with appropriate antibiotics, serially diluted, spotted on LB plates with antibiotics, with/without arabinose, and then incubated at 37°C. Representative data shown from two independent experiments.

## SUPPLEMENTARY INFORMATION

**Supplementary Table 1.** Proteomics data of RT-3xFlag/RcaT-3xFlag pull-downs.

**Supplementary Table 2.** Genotypes of bacterial strains used in this study.

**Supplementary Table 3.** Description of plasmids used in this study.

**Supplementary Table 4.** Description of construction of plasmids used in this study.

**Supplementary Table 5.** List of primers used in this study.

