## Supplementary Table 2. Genotypes of bacterial strains used in this study. for "Bacterial retrons encode tripartite toxin/antitoxin systems"

**Supplementary Table 2.** Genotypes of bacterial strains used in this study (1/6).

| **Name (as in the figures)** | **Genotype** | **Strain code** | **Used in** | **Used for** |
|---|---|---|---|---|
| WT | *Salmonella enterica* enterica Typhimurium 14028s | *S*Tm | Fig. 1D, ED Fig. 1A-1C | Cold-sensitivity, anaerobic growth assay |
| Δ*rrtT* | *S*Tm ΔSTM14_4641:FRT (Δ*rrtT*::FRT) | NT16019 | Fig. 1D, ED Fig. 1A-1C | Cold-sensitivity, anaerobic growth assay |
| Δ*xseA* | *S*Tm Δ*xseA*::FRT | NT16190 | Fig. 1D, ED Fig. 1A-1C | Cold-sensitivity, anaerobic growth assay |
| Δ*xseB* | *S*Tm Δ*xseB*::FRT | NT16487 | Fig. 1D, ED Fig. 1A-1C | Cold-sensitivity, anaerobic growth assay |
| Δ*rnhA* | *S*Tm Δ*rnhA*::FRT | NT16276 | Fig. 1D, ED Fig. 1A-1C | Cold-sensitivity, anaerobic growth assay |
| Δ*msrmsd* | *S*Tm Δ*msrmsd*::FRT | NT16085 | Fig. 1D, ED Fig. 1A-1C | Cold-sensitivity, anaerobic growth assay |
| ΔSTM14_4640 | *S*Tm ΔSTM14_4640::FRT (Δ*rcaT*::FRT) | NT16277 | Fig. 1D, ED Fig. 1A-1C | Cold-sensitivity, anaerobic growth assay |
| WT | *Salmonella enterica* enterica Typhimurium 14028s | *S*Tm | Fig. 2B/2C | Cold-sensitivity/anaerobic growth assay |
| Δ*rcaT* | *S*Tm Δ*rcaT*::FRT | NT16277 | Fig. 2B/2C | Cold-sensitivity/anaerobic growth assay |
| Δ*xseA* | *S*Tm Δ*xseA*::kan | NT16018 | Fig. 2B/2C | Cold-sensitivity/anaerobic growth assay |
| Δ*xseA* Δ*rcaT* | *S*Tm Δ*rcaT*::FRT ΔxseA::kan | NT16219 | Fig. 2B/2C | Cold-sensitivity/anaerobic growth assay |
| Δ*xseB* | *S*Tm Δ*xseB*::kan | NT16484 | Fig. 2B/2C | Cold-sensitivity/anaerobic growth assay |
| Δ*xseB* Δ*rcaT* | *S*Tm Δ*rcaT*::FRT Δ*xseB*::kan | NT16497 | Fig. 2B/2C | Cold-sensitivity/anaerobic growth assay |
| Δ*rnhA* | *S*Tm Δ*rnhA*::kan | NT16245 | Fig. 2B/2C | Cold-sensitivity/anaerobic growth assay |
| Δ*rnhA* Δ*rcaT* | *S*Tm Δ*rcaT*::FRT Δ*rnhA*::kan | NT16213 | Fig. 2B/2C | Cold-sensitivity/anaerobic growth assay |
| p-empty | *E. coli* BL21-AI DE3 pET28α | NT16297 | Fig. 2D | RcaT over-expression growth assay |
| p-*rcaT* | *E. coli* BL21-AI DE pJB11 | NT16294 | Fig. 2D | RcaT over-expression growth assay |
| p-*rcaT*$^{D296V}$ | *E. coli* BL21-AI DE3 pJB24 | NT16295 | Fig. 2D | RcaT over-expression growth assay |
| WT | *Salmonella enterica* enterica Typhimurium 14028s | *S*Tm | Fig. 3A/3B, ED Fig. 6A, ED Fig. 8A | Immunoprecipitation (control for *rrtT*-3xFlag strains) |
| WT *rrtT*-3xFlag | *S*Tm *rrtT*-2xStrep-TEV-3xFLAG:FRT | NT16869 | Fig. 3A/3B, ED Fig. 6A, ED Fig. 8A | Immunoprecipitation |
| Δ*xseA* *rrtT*-3xFlag | *S*Tm *rrtT*-2xStrep-TEV-3xFlag:FRT Δ*xseA*::kan | NT16969 | Fig. 3A/3B, ED Fig. 6A, ED Fig. 8A | Immunoprecipitation |
| Δ*msrmsd* *rrtT*-3xFlag | *S*Tm *rrtT*-2xStrep-TEV-3xFlag:FRT Δ*msrmsd*::kan | NT16938 | Fig. 3A/3B, ED Fig. 6A, ED Fig. 8A | Immunoprecipitation |
| ΔSTM14_4645 | *S*Tm ΔSTM14_4645::cat | NT16784 | Fig. 3A/3B, ED Fig. 6A, ED Fig. 8A | Immunoprecipitation (control for *rcaT*-3xFlag strains) |
| WT *rcaT*-3xFlag | *S*Tm ΔSTM14_4645::cat *rcaT*-3xFlag | NT16939 | Fig. 3C/3D, ED Fig. 6B, ED Fig. 8A | Immunoprecipitation |
| Δ*xseA* *rcaT*-3xFlag | *S*Tm ΔxseA::FRT ΔSTM14_4645::cat *rcaT*-3xFlag | NT16945 | Fig. 3C/3D, ED Fig. 6B, ED Fig. 8A | Immunoprecipitation |
| Δ*msd* *rcaT*-3xFlag | *S*Tm ΔSTM14_4645::cat Δ79:*msd* *rcaT*-3xFlag | NT16944 | Fig. 3C/3D, ED Fig. 6B, ED Fig. 8A | Immunoprecipitation |
| $P_{BAD}$-RT (Se) $P_{tac}$-msrmsd (-) | *E. coli* BW25113 pJB141 pNTR-SD | NT32230 | Fig. 4A | Hybrid-retron msDNA isolation |
| $P_{BAD}$-RT (Se) $P_{tac}$-msrmsd (Se) | *E. coli* BW25113 pJB141 pKM1 | NT32236 | Fig. 4A | Hybrid-retron msDNA isolation |
| $P_{BAD}$-RT (Se) $P_{tac}$-msrmsd (Ec) | *E. coli* BW25113 pJB141 pJB137 | NT32243 | Fig. 4A | Hybrid-retron msDNA isolation |
| $P_{BAD}$-RT (Ec) $P_{tac}$-msrmsd (-) | *E. coli* BW25113 pJB142 pNTR-SD | NT32231 | Fig. 4A | Hybrid-retron msDNA isolation |
| $P_{BAD}$-RT (Ec) $P_{tac}$-msrmsd (Se) | *E. coli* BW25113 pJB142 pKM1 | NT32238 | Fig. 4A | Hybrid-retron msDNA isolation |

**Supplementary Table 2.** Genotypes of bacterial strains used in this study (2/6).

| | | | | |
|---|---|---|---|---|
| $P_{BAD}$-RT (Ec) $P_{tac}$-msrmsd (Ec) | *E. coli* BW25113 pJB142 pJB137 | NT32244 | Fig. 4A | Hybrid-retron msDNA isolation |
| $P_{BAD}$-RcaT-RT (Se-Se) $P_{tac}$-msrmsd (-) | *E. coli* BW25113 pJB87 pNTR-SD | NT32226 | Fig. 4B/ED Fig. 10F | Hybrid-retron growth assay |
| $P_{BAD}$-RcaT-RT (Se-Se) $P_{tac}$-msrmsd (Se) | *E. coli* BW25113 pJB87 pKM1 | NT32233 | Fig. 4B/ED Fig. 10F | Hybrid-retron growth assay |
| $P_{BAD}$-RcaT-RT (Se-Se) $P_{tac}$-msrmsd (Ec) | *E. coli* BW25113 pJB87 pJB137 | NT32239 | Fig. 4B/ED Fig. 10F | Hybrid-retron growth assay |
| $P_{BAD}$-RcaT-RT (Se-Ec) $P_{tac}$-msrmsd (-) | *E. coli* BW25113 pJB139 pNTR-SD | NT32228 | Fig. 4B/ED Fig. 10F | Hybrid-retron growth assay |
| $P_{BAD}$-RcaT-RT (Se-Ec) $P_{tac}$-msrmsd (Se) | *E. coli* BW25113 pJB139 pKM1 | NT32235 | Fig. 4B/ED Fig. 10F | Hybrid-retron growth assay |
| $P_{BAD}$-RcaT-RT (Se-Ec) $P_{tac}$-msrmsd (Ec) | *E. coli* BW25113 pJB139 pJB137 | NT32241 | Fig. 4B/ED Fig. 10F | Hybrid-retron growth assay |
| $P_{BAD}$-RcaT-RT (Ec-Se) $P_{tac}$-msrmsd (-) | *E. coli* BW25113 pJB138 pNTR-SD | NT32227 | Fig. 4B/ED Fig. 10F | Hybrid-retron growth assay |
| $P_{BAD}$-RcaT-RT (Ec-Se) $P_{tac}$-msrmsd (Se) | *E. coli* BW25113 pJB138 pKM1 | NT32234 | Fig. 4B/ED Fig. 10F | Hybrid-retron growth assay |
| $P_{BAD}$-RcaT-RT (Ec-Se) $P_{tac}$-msrmsd (Ec) | *E. coli* BW25113 pJB138 pJB137 | NT32240 | Fig. 4B/ED Fig. 10F | Hybrid-retron growth assay |
| $P_{BAD}$-RcaT-RT (Ec-Ec) $P_{tac}$-msrmsd (-) | *E. coli* BW25113 pJB140 pNTR-SD | NT32229 | Fig. 4B/ED Fig. 10F | Hybrid-retron growth assay |
| $P_{BAD}$-RcaT-RT (Ec-Ec) $P_{tac}$-msrmsd (Se) | *E. coli* BW25113 pJB140 pKM1 | NT32236 | Fig. 4B/ED Fig. 10F | Hybrid-retron growth assay |
| $P_{BAD}$-RcaT-RT (Ec-Ec) $P_{tac}$-msrmsd (Ec) | *E. coli* BW25113 pJB140 pJB137 | NT32242 | Fig. 4B/ED Fig. 10F | Hybrid-retron growth assay |
| WT | *S* Tm Δ*araBAD* ::kan pJB51 | NT16732 | ED Fig. 1D | msDNA-Sen2 isolation |
| Δ*rnhA* | *S* Tm Δ*rnhA* ::FRT Δ*araBAD* ::kan pJB51 | NT16733 | ED Fig. 1D | msDNA-Sen2 isolation |
| Δ*xseA* | *S* Tm Δ*xseA* ::FRT Δ*araBAD* ::kan pJB51 | NT16659 | ED Fig. 1D | msDNA-Sen2 isolation |
| Δ*xseB* | *STm* Δ*xseB* ::FRT Δ*araBAD* ::kan pJB51 | NT16734 | ED Fig. 1D | msDNA-Sen2 isolation |
| ΔSTM14_4640 | *STm* Δ*rcaT* ::FRT Δ*araBAD* ::kan pJB51 | NT16735 | ED Fig. 1D | msDNA-Sen2 isolation |
| WT | *Salmonella enterica* enterica Typhimurium 14028s | *S* Tm | ED Fig. 2A | Cold-sensitivity growth assay |
| Δ*rrtT* Suppressor 1 | *S* Tm Δ*rrtT* ::FRT *rcaT* -(TTTA)$_{2\rightarrow3}$ (417/963) | NT16067 | ED Fig. 2A | Cold-sensitivity growth assay |
| Δ*rrtT* Suppressor 2 | *S* Tm Δ*rrtT* ::FRT *rcaT* -A$_{7\rightarrow6}$ (361/963) | NT16068 | ED Fig. 2A | Cold-sensitivity growth assay |
| Δ*rrtT* Suppressor 3 | *S* Tm Δ*rrtT* ::FRT *rcaT* -S152T (TCT→ACT) | NT16069 | ED Fig. 2A | Cold-sensitivity growth assay |
| Δ*rrtT* Suppressor 4 | *S* Tm Δ*rrtT* ::FRT *rcaT* -S253G (AGC→GGC) | NT16070 | ED Fig. 2A | Cold-sensitivity growth assay |
| Δ*rrtT* Suppressor 5 | *S* Tm Δ*rrtT* ::FRT *trmE*→*rcaT* -Δ62 (-100) | NT16074 | ED Fig. 2A | Cold-sensitivity growth assay |
| Δ*rrtT* Suppressor 6 | *S* Tm Δ*rrtT* ::FRT *rcaT* -D296Y (GAT→TAT) | NT16075 | ED Fig. 2A | Cold-sensitivity growth assay |
| Δ*rrtT* Suppressor 7 | *S* Tm Δ*rrtT* ::FRT *rcaT* -(TTTA)$_{2\rightarrow3}$ (417/963) | NT16076 | ED Fig. 2A | Cold-sensitivity growth assay |
| Δ*rrtT* Suppressor 8 | *S* Tm Δ*rrtT* ::FRT *rcaT* -Δ88 (-77/+11) | NT16077 | ED Fig. 2A | Cold-sensitivity growth assay |
| Δ*xseA* | *S* Tm Δ*xseA* ::kan | NT16018 | ED Fig. 2A | Cold-sensitivity growth assay |
| Δ*xseA* Suppressor 1 | STm Δ*xseA* ::kan *rcaT* -(TTTA)$_{2\rightarrow3}$ (417/963) | NT16160 | ED Fig. 2A | Cold-sensitivity growth assay |
| Δ*xseA* Suppressor 2 | STm Δ*xseA* ::kan *rcaT* -(TTTA)$_{2\rightarrow3}$ (417/963) | NT16163 | ED Fig. 2A | Cold-sensitivity growth assay |
| Δ*xseA* Suppressor 3 | STm Δ*xseA* ::kan *rcaT* -D296V (GAT→GTT) | NT16164 | ED Fig. 2A | Cold-sensitivity growth assay |
| Δ*xseA* Suppressor 4 | *S* Tm Δ*xseA* ::kan *rcaT* -Δ12 (510-521/963) | NT16071 | ED Fig. 2A | Cold-sensitivity growth assay |

**Supplementary Table 2.** Genotypes of bacterial strains used in this study (3/6).

| | | | | |
|---|---|---|---|---|
| Δ*xseA* Suppressor 5 | *S*Tm Δ*xseA*::kan *rcaT*-E180* (GAA→TAA) | NT16072 | ED Fig. 2A | Cold-sensitivity growth assay |
| Δ*xseA* Suppressor 6 | *S*Tm Δ*xseA*::kan *rcaT*-T7→6 (296/963) | NT16073 | ED Fig. 2A | Cold-sensitivity growth assay |
| Δ*xseA* Suppressor 7 | *S*Tm Δ*xseA*::kan *rcaT*-(TTTA)2→3 (417/963) | NT16078 | ED Fig. 2A | Cold-sensitivity growth assay |
| Δ*xseA* Suppressor 8 | *S*Tm Δ*xseA*::kan *rcaT*-E180* (GAA→TAA) | NT16079 | ED Fig. 2A | Cold-sensitivity growth assay |
| Δ*msrmsd* | *S*Tm Δ*msrmsd*::kan | NT16082 | ED Fig. 2A | Cold-sensitivity growth assay |
| Δ*msrmsd* Suppressor 1 | *S*Tm Δ*msrmsd*::FRT *rcaT*-C6→5 (126/963) | NT16122 | ED Fig. 2A | Cold-sensitivity growth assay |
| Δ*msrmsd* Suppressor 2 | *S*Tm Δ*msrmsd*::FRT *rcaT*-D160V (GAT→GTT) | NT16131 | ED Fig. 2A | Cold-sensitivity growth assay |
| Δ*msrmsd* Suppressor 3 | *S*Tm Δ*msrmsd*::FRT *rcaT*-(TTTA)2→3 (417/963) | NT16148 | ED Fig. 2A | Cold-sensitivity growth assay |
| Δ*msrmsd* Suppressor 4 | *S*Tm Δ*msrmsd*::FRT *rcaT*-A5→6 (697/963) | NT16149 | ED Fig. 2A | Cold-sensitivity growth assay |
| Δ*msrmsd* Suppressor 5 | *S*Tm Δ*msrmsd*::FRT *rcaT*-D296V (GAT→GTT) | NT16150 | ED Fig. 2A | Cold-sensitivity growth assay |
| Δ*msrmsd* Suppressor 6 | *S*Tm Δ*msrmsd*::FRT *rcaT*-D160G (GAT→GGT) | NT16152 | ED Fig. 2A | Cold-sensitivity growth assay |
| Δ*msrmsd* Suppressor 7 | *S*Tm Δ*msrmsd*::FRT *rcaT*-Δ10 (356-265/963) | NT16153 | ED Fig. 2A | Cold-sensitivity growth assay |
| Δ*msrmsd* Suppressor 8 | *S*Tm Δ*msrmsd*::FRT *rcaT*-E180* (GAA→TAA) | NT16154 | ED Fig. 2A | Cold-sensitivity growth assay |
| Δ*msrmsd* Suppressor 9 | *S*Tm Δ*msrmsd*::FRT *rcaT*-E217* (GAA→TAA) | NT16155 | ED Fig. 2A | Cold-sensitivity growth assay |
| Δ*msrmsd* Suppressor 10 | *S*Tm Δ*msrmsd*::FRT *rcaT*-D160V (GAT→GTT) | NT16156 | ED Fig. 2A | Cold-sensitivity growth assay |
| Δ*msrmsd* Suppressor 11 | *S*Tm Δ*msrmsd*::FRT *rcaT*-A7→6 (361/963) | NT16157 | ED Fig. 2A | Cold-sensitivity growth assay |
| Δ*msrmsd* Suppressor 12 | *S*Tm Δ*msrmsd*::FRT *rcaT*-D296V (GAT→GTT) | NT16158 | ED Fig. 2A | Cold-sensitivity growth assay |
| Δ*msrmsd* Suppressor 13 | *S*Tm Δ*msrmsd*::FRT *rcaT*-G197V (GGA→GTA) | NT16159 | ED Fig. 2A | Cold-sensitivity growth assay |
| WT intact | *Salmonella enterica* enterica Typhimurium 14028s | *S*Tm | ED Fig. 2C/2D | Cold-sensitivity growth assay/Western blot |
| WT *msrmsd* | *S*Tm ΔSTM14_4645::cat Δ0:*msd* *rcaT*-3xFlag | NT16939 | ED Fig. 2C/2D | Cold-sensitivity growth assay/Western blot |
| WT Δ62:*msd* | *S*Tm ΔSTM14_4645::cat Δ62:*msd* *rcaT*-3xFlag | NT16940 | ED Fig. 2C/2D | Cold-sensitivity growth assay/Western blot |
| WT Δ63:*msd* | *S*Tm ΔSTM14_4645::cat Δ63:*msd* *rcaT*-3xFlag | NT16941 | ED Fig. 2C/2D | Cold-sensitivity growth assay/Western blot |
| WT Δ67:*msd* | *S*Tm ΔSTM14_4645::cat Δ67:*msd* *rcaT*-3xFlag | NT16942 | ED Fig. 2C/2D | Cold-sensitivity growth assay/Western blot |
| WT Δ71:*msd* | *S*Tm ΔSTM14_4645::cat Δ71:*msd* *rcaT*-3xFlag | NT16943 | ED Fig. 2C/2D | Cold-sensitivity growth assay/Western blot |
| WT Δ79:*msd* | *S*Tm ΔSTM14_4645::cat Δ79:*msd* *rcaT*-3xFlag | NT16944 | ED Fig. 2C/2D | Cold-sensitivity growth assay/Western blot |
| Δ*xseA* intact | *S*Tm Δ*xseA*::FRT | NT16190 | ED Fig. 2C | Cold-sensitivity growth assay |
| ΔxseA msrmsd | *S*Tm Δ*xseA*::FRT ΔSTM14_4645::cat Δ0:*msd* *rcaT*-3xFlag | NT16945 | ED Fig. 2C | Cold-sensitivity growth assay |
| Δ*xseA* Δ62:msd | *S*Tm Δ*xseA*::FRT ΔSTM14_4645::cat Δ62:*msd* *rcaT*-3xFlag | NT16946 | ED Fig. 2C | Cold-sensitivity growth assay |
| Δ*xseA* Δ63:msd | *S*Tm Δ*xseA*::FRT ΔSTM14_4645::cat Δ63:*msd* *rcaT*-3xFlag | NT16947 | ED Fig. 2C | Cold-sensitivity growth assay |
| Δ*xseA* Δ67:msd | *STm ΔxseA*::FRT ΔSTM14_4645::cat Δ67:*msd* *rcaT*-3xFlag | NT16948 | ED Fig. 2C | Cold-sensitivity growth assay |
| Δ*xseA* Δ71:msd | *S*Tm Δ*xseA*::FRT ΔSTM14_4645::cat Δ71:*msd* *rcaT*-3xFlag | NT16949 | ED Fig. 2C | Cold-sensitivity growth assay |
| Δ*xseA* Δ79:msd | *S*Tm Δ*xseA*::FRT ΔSTM14_4645::cat Δ79:*msd* *rcaT*-3xFlag | NT16950 | ED Fig. 2C | Cold-sensitivity growth assay |

**Supplementary Table 2.** Genotypes of bacterial strains used in this study (4/6).

| | | | | |
|---|---|---|---|---|
| WT | *Salmonella enterica* enterica Typhimurium 14028s | *S*Tm | ED Fig. 3A/3B | Native RcaT toxicity assay (growth rate/viability) |
| Δ*msrmsd* | *S*Tm Δ*msrmsd*::FRT | NT16085 | ED Fig. 3A/3B | Native RcaT toxicity assay (growth rate/viability) |
| p-empty | *E. coli* BW25113 pBAD30 | NT16335 | ED Fig. 3C/3D | Over-expression RcaT toxicity assay (growth rate/viability) |
| p-*rcaT* | *E. coli* BW25113 pJB12 | NT16336 | ED Fig. 3C/3D | Over-expression RcaT toxicity assay (growth rate/viability) |
| p-empty | *E. coli* BW25113 pTU175 | NT16799 | ED Fig. 4A | RcaT over-expression growth assay |
| p-*rcaT* | *E. coli* BW25113 pJB37 | NT16593 | ED Fig. 4A | RcaT over-expression growth assay |
| p-retron | *E. coli* BW25113 pJB38 | NT16585 | ED Fig. 4A | RcaT over-expression growth assay |
| p-retron-Δ*rrtT* | *E. coli* BW25113 pJB45 | NT16961 | ED Fig. 4A | RcaT over-expression growth assay |
| p-retron-Δ*msrmsd* | *E. coli* BW25113 pJB87 | NT16962 | ED Fig. 4A | RcaT over-expression growth assay |
| p-retron-*msrmsd*$^{mut}$ | *E. coli* BW25113 pJB86 | NT16957 | ED Fig. 4A | RcaT over-expression growth assay |
| p-retron-Δ*rcaT* | *E. coli* BW25113 pJB51 | NT16651 | ED Fig. 4A | RcaT over-expression growth assay |
| p-empty + p-empty | *E. coli* BW25113 pBAD33 pTU175 | NT16963 | ED Fig. 4B | RcaT over-expression growth assay |
| p-empty + p-retron-Δ*rcaT* | *E. coli* BW25113 pBAD33 pJB51 | NT16964 | ED Fig. 4B | RcaT over-expression growth assay |
| p-*rcaT* + p-empty | *E. coli* BW25113 pJB15 pTU175 | NT16965 | ED Fig. 4B | RcaT over-expression growth assay |
| p-*rcaT* + p-retron-Δ*rcaT* | *E. coli* BW25113 pJB15 pJB51 | NT16966 | ED Fig. 4B | RcaT over-expression growth assay |
| WT p-retron-Δ*rcaT* | *E. coli* BW25113 pJB51 | NT16651 | ED Fig. 4C | RcaT over-expression growth assay |
| WT p-retron | *E. coli* BW25113 pJB38 | NT16585 | ED Fig. 4C | RcaT over-expression growth assay |
| Δ*xseA* p-retron-Δ*rcaT* | *E. coli* BW25113 Δ*xseA*::kan pJB51 | NT32248 | ED Fig. 4C | RcaT over-expression growth assay |
| Δ*xseA* p-retron | *E. coli* BW25113 Δ*xseA*::kan pJB38 | NT32245 | ED Fig. 4C | RcaT over-expression growth assay |
| Δ*xseB* p-retron-Δ*rcaT* | *E. coli* BW25113 Δ*xseB*::kan pJB51 | NT32249 | ED Fig. 4C | RcaT over-expression growth assay |
| Δ*xseB* p-retron | *E. coli* BW25113 Δ*xseB*::kan pJB38 | NT32246 | ED Fig. 4C | RcaT over-expression growth assay |
| Δ*rnhA* p-retron-Δ*rcaT* | *E. coli* BW25113 Δ*rnhA*::kan pJB51 | NT32250 | ED Fig. 4C | RcaT over-expression growth assay |
| Δ*rhnA* p-retron | *E. coli* BW25113 Δ*rnhA*::kan pJB38 | NT32247 | ED Fig. 4C | RcaT over-expression growth assay |
| Native | *Salmonella enterica* enterica Typhimurium 14028s | *S*Tm | ED Fig. 5A | Western blot (control) |
| WT *rcaT*-3xFlag | *S*Tm ΔSTM14_4645::cat *rcaT*-3xFlag | NT16939 | ED Fig. 5A/5B | Western blot |
| Δ*rrtT* *rcaT*-3xFlag | *S*Tm ΔSTM14_4645::cat Δ*rrtT*::kan *rcaT*-3xFlag | NT16936 | ED Fig. 5A/5B | Western blot |
| Δ*msd* *rcaT*-3xFlag | *S*Tm ΔSTM14_4645::cat Δ79:*msd* *rcaT*-3xFlag | NT16944 | ED Fig. 5A/5B | Western blot |
| Δ*xseA* *rcaT*-3xFlag | *S*Tm Δ*xseA*::FRT ΔSTM14_4645::cat *rcaT*-3xFlag | NT16812 | ED Fig. 5A/5B | Western blot |
| Δ*rnhA* *rcaT*-3xFlag | *S*Tm Δ*rnhA*::FRT ΔSTM14_4645::cat *rcaT*-3xFlag | NT16813 | ED Fig. 5A/5B | Western blot |
| WT | *Salmonella enterica* enterica Typhimurium 14028s | *S*Tm | ED Fig. 5C | Cold-sensitivity growth assay |
| WT* | *S*Tm ΔSTM14_4645::cat | NT16784 | ED Fig. 5C | Cold-sensitivity growth assay |
| WT* *rcaT*-3xFlag | *S*Tm ΔSTM14_4645::cat *rcaT*-3xFlag | NT16939 | ED Fig. 5C | Cold-sensitivity growth assay |

**Supplementary Table 2.** Genotypes of bacterial strains used in this study (5/6).

| | | | | |
|---|---|---|---|---|
| Δ*rrtT* | *S*Tm Δ*rrtT*::kan | NT16000 | ED Fig. 5C | Cold-sensitivity growth assay |
| Δ*rrtT* * *rcaT*-3xFlag | *S*Tm ΔSTM14_4645::cat Δ*rrtT*::kan *rcaT*-3xFlag | NT16936 | ED Fig. 5C | Cold-sensitivity growth assay |
| Δ*xseA* | *S*Tm Δ*xseA*::kan | NT16018 | ED Fig. 5C | Cold-sensitivity growth assay |
| Δ*xseA* * *rcaT*-3xFlag | *S*Tm Δ*xseA*::FRT ΔSTM14_4645::cat *rcaT*-3xFlag | NT16812 | ED Fig. 5C | Cold-sensitivity growth assay |
| Δ*rnhA* | *S*Tm Δ*rnhA*::kan | NT16245 | ED Fig. 5C | Cold-sensitivity growth assay |
| Δ*rnhA* * *rcaT*-3xFlag | *S*Tm Δ*rnhA*::FRT ΔSTM14_4645::cat *rcaT*-3xFlag | NT16813 | ED Fig. 5C | Cold-sensitivity growth assay |
| Δ*msrmsd* | *S*Tm Δ*msrmsd*::kan | NT16082 | ED Fig. 5C | Cold-sensitivity growth assay |
| Δ*msrmsd* * *rcaT*-3xFlag | *S*Tm ΔSTM14_4645::cat Δ*msrmsd*::kan *rcaT*-3xFlag | NT16937 | ED Fig. 5C | Cold-sensitivity growth assay |
| WT | *Salmonella enterica* enterica Typhimurium 14028s | *S*Tm | ED Fig. 7 | Cold-sensitivity growth assay |
| *rrtT*-3xFlag | *S*Tm *rrtT*-2xStrep-TEV-3xFlag:FRT | NT16869 | ED Fig. 7 | Cold-sensitivity growth assay |
| Δ*xseA* | *S*Tm Δ*xseA*::kan | NT16018 | ED Fig. 7 | Cold-sensitivity growth assay |
| Δ*xseA* *rrtT*-3xFlag | *S*Tm *rrtT*-2xStrep-TEV-3xFlag:FRT Δ*xseA*::kan | NT16969 | ED Fig. 7 | Cold-sensitivity growth assay |
| Δ*rnhA* | *S*Tm Δ*rnhA*::kan | NT16245 | ED Fig. 7 | Cold-sensitivity growth assay |
| Δ*rnhA* *rrtT*-3xFlag | *S*Tm *rrtT*-2xStrep-TEV-3xFlag:FRT Δ*rnhA*::kan | NT16970 | ED Fig. 7 | Cold-sensitivity growth assay |
| Δ*msrmsd* | *S*Tm Δ*msrmsd*::kan | NT16082 | ED Fig. 7 | Cold-sensitivity growth assay |
| Δ*msrmsd* *rrtT*-3xFlag | *S*Tm *rrtT*-2xStrep-TEV-3xFlag:FRT Δ*msrmsd*::kan | NT16875 | ED Fig. 7 | Cold-sensitivity growth assay |
| RT-Sen2-6xHis | *E. coli* BL21 DE3 CodonPlus-RIL pJB120 | NT32223 | ED Fig. 9A/9C | RT-Sen2-His purification/DNA isolation from purified RT |
| p-*rcaT* p-empty | *E. coli* BL21 DE3 AI pET28 pJB37 | NT32151 | ED Fig. 9B | RcaT over-expression growth assay |
| p-*rcaT* p-*msrmsd*-*rrtT*-6xHis | *E. coli* BL21 DE3 AI pJB120 pJB37 | NT32152 | ED Fig. 9B | RcaT over-expression growth assay |
| p-empty | *E. coli* BW25113 pBAD33 | NT16428 | ED Fig. 10C | RcaT-Eco9 over-expression growth assay |
| p-rcaT-Eco9 | *E. coli* BW25113 pJB91 | NT32018 | ED Fig. 10C | RcaT-Eco9 over-expression growth assay |
| p-retron-Eco9 | *E. coli* BW25113 pJB92 | NT32019 | ED Fig. 10C | RcaT-Eco9 over-expression growth assay |
| WT Ptac-msrmsd-Eco9 (+) PBAD-RT-Eco9 (+) PBAD-RcaT-RT-Eco9 (-) | *E. coli* BW25113 pJB137 pJB142 | NT32255 | ED Fig. 10D | Retron-Eco9 over-expression growth assay |
| WT Ptac-msrmsd-Eco9 (+) PBAD-RT-Eco9 (-) PBAD-RcaT-RT-Eco9 (+) | *E. coli* BW25113 pJB137 pJB140 | NT32251 | ED Fig. 10D | Retron-Eco9 over-expression growth assay |
| ΔxseA Ptac-msrmsd-Eco9 (+) PBAD-RT-Eco9 (+) PBAD-RcaT-RT-Eco9 (-) | *E. coli* BW25113 Δ*xseA* pJB137 pJB142 | NT32256 | ED Fig. 10D | Retron-Eco9 over-expression growth assay |
| ΔxseA Ptac-msrmsd-Eco9 (+) PBAD-RT-Eco9 (-) PBAD-RcaT-RT-Eco9 (+) | *E. coli* BW25113 Δ*xseA* pJB137 pJB140 | NT32252 | ED Fig. 10D | Retron-Eco9 over-expression growth assay |
| ΔxseB Ptac-msrmsd-Eco9 (+) PBAD-RT-Eco9 (+) PBAD-RcaT-RT-Eco9 (-) | *E. coli* BW25113 Δ*xseB* pJB137 pJB142 | NT32257 | ED Fig. 10D | Retron-Eco9 over-expression growth assay |
| ΔxseB Ptac-msrmsd-Eco9 (+) PBAD-RT-Eco9 (-) PBAD-RcaT-RT-Eco9 (+) | *E. coli* BW25113 Δ*xseB* pJB137 pJB140 | NT32253 | ED Fig. 10D | Retron-Eco9 over-expression growth assay |
| ΔrnhA Ptac-msrmsd-Eco9 (+) PBAD-RT-Eco9 (+) PBAD-RcaT-RT-Eco9 (-) | *E. coli* BW25113 Δ*rnhA* pJB137 pJB142 | NT32258 | ED Fig. 10D | Retron-Eco9 over-expression growth assay |
| ΔrnhA Ptac-msrmsd-Eco9 (+) PBAD-RT-Eco9 (-) PBAD-RcaT-RT-Eco9 (+) | *E. coli* BW25113 Δ*rnhA* pJB137 pJB140 | NT32254 | ED Fig. 10D | Retron-Eco9 over-expression growth assay |
| WT PBAD-msrmsd-RT-Eco9 | *E. coli* BW25113 pJB95 | NT32060 | ED Fig. 10F | msDNA-Eco9 isolation |
| ΔxseA PBAD-msrmsd-RT-Eco9 | *E. coli* BW25113 Δ*xseA*::kan pJB95 | NT32063 | ED Fig. 10F | msDNA-Eco9 isolation |

**Supplementary Table 2.** Genotypes of bacterial strains used in this study (6/6).

| | | | | |
|---|---|---|---|---|
| ΔxseB PBAD-msrmsd-RT-Eco9 | *E. coli* BW25113 Δ*xseB*::kan pJB95 | NT32064 | ED Fig. 10F | msDNA-Eco9 isolation |
| ΔrnhA PBAD-msrmsd-RT-Eco9 | *E. coli* BW25113 Δ*rnhA*::kan pJB95 | NT32065 | ED Fig. 10F | msDNA-Eco9 isolation |
| Δ63 Δ*rrtT* suppressor | *S*Tm Δ*rrtT*::FRT Δ63:*msd* | NT16489 | - | Construction of NT16941 |
