## Supplementary Table 3. Description of plasmids used in this study. for "Bacterial retrons encode tripartite toxin/antitoxin systems"

**Supplementary Table 3.** Description of plasmids used in this study (1/1).

| Plasmid code | Plasmid name | Backbone | Used for | Antibiotic resistance | Reference |
|---|---|---|---|---|---|
| pJB11 | pET28α-His-*rcaT* | pET28α | Over-expression of His-tagged-*rcaT*. | Kanamycin | This work |
| pJB12 | pBAD30-*rcaT* | pBAD30 | Over-expression of *rcaT*. | Ampicillin | This work |
| pJB120 | pET28α-*msrmsd*-*rrtT*-TEV-His | pET28α | Over-expression of *msrmsd*-*rrtT*-TEV-His. | Kanamycin | This work |
| pJB15 | pBAD33-*rcaT* | pBAD33 | Over-expression of *rcaT*. | Chloramphenicol | This work |
| pJB24 | pET28α-His-*rcaT*-D296V | pET28α | Over-expression of His-*rcaT*-D296V. | Kanamycin | This work |
| pJB37 | pTU175-pBAD-*rcaT* | pTU175 | Over-expression of *rcaT*. | Spectinomycin | This work |
| pJB38 | pTU175-pBAD-retron | pTU175 | Over-expression of retron. | Spectinomycin | This work |
| pJB45 | pTU175-pBAD-*msrmsd*-*rcaT* | pTU175 | Over-expression of *msrmsd*-*rcaT*. | Spectinomycin | This work |
| pJB46 | pTU175-pBAD-Δ62*msd*-*rcaT* | pTU175 | Construction of NT16940, NT16946. | Spectinomycin | This work |
| pJB47 | pTU175-pBAD-Δ63*msd*-*rcaT* | pTU175 | Construction of NT16941, NT16947. | Spectinomycin | This work |
| pJB48 | pTU175-pBAD-Δ71*msd*-*rcaT* | pTU175 | Construction of NT16943, NT16949. | Spectinomycin | This work |
| pJB49 | pTU175-pBAD-Δ79*msd*-*rcaT* | pTU175 | Construction of NT16944, NT16950. | Spectinomycin | This work |
| pJB51 | pTU175-*msrmsd*-*rrtT* (RT-Sen2) | pTU175 | Over-expression of *msrmsd*-*rrtT*. | Spectinomycin | This work |
| pJB64 | pTU175-pBAD-Δ67*msd*-*rcaT* | pTU175 | Construction of NT16942, NT16948. | Spectinomycin | This work |
| pJB86 | pJB38-bG→T-retron | pTU175 | Over-expression of *msr* bG→T-retron. | Spectinomycin | This work |
| pJB87 | pJB37-*rcaT*-*rrtT* | pTU175 | Over-expression of *rcaT*-*rrtT*. | Spectinomycin | This work |
| pJB91 | pBAD33-rcaT(Eco9) | pBAD33 | Over-expression of rcaT(Eco9). | Chloramphenicol | This work |
| pJB92 | pBAD33-retron(Eco9) | pBAD33 | Over-expression of retron(Eco9). | Chloramphenicol | This work |
| pJB93 | pTU175-pBAD-msrmsd(Eco9)-RT(Sen2) | pTU175 | Over-expression of msrmsd(Eco9)-rrtT. | Spectinomycin | This work |
| pJB94 | pTU175-pBAD-msrmsd(Sen2)-RT(Eco9) | pTU175 | Over-expression of msrmsd(Sen2)-RT(Eco9). | Spectinomycin | This work |
| pJB95 | pTU175-pBAD-msrmsd(Eco9)-RT(Eco9) | pTU175 | Over-expression of msrmsd(Eco9)-RT(Eco9). | Spectinomycin | This work |
| pJB138 | pTU175-pBAD-rcaT(Eco9)-RT(Sen2) | pTU175 | Over-expression of rcaT(Eco9)-RT(Sen2). | Spectinomycin | This work |
| pJB139 | pTU175-pBAD-rcaT(Sen2)-RT(Eco9) | pTU175 | Over-expression of rcaT(Sen2)-RT(Eco9). | Spectinomycin | This work |
| pJB140 | pTU175-pBAD-rcaT(Eco9)-RT(Eco9) | pTU175 | Over-expression of rcaT(Eco9)-RT(Eco9). | Spectinomycin | This work |
| pJB141 | pTU175-pBAD-RT(Sen2) | pTU175 | Over-expression of RT(Sen2). | Spectinomycin | This work |
| pJB142 | pTU175-pBAD-RT(Eco9) | pTU175 | Over-expression of RT(Eco9). | Spectinomycin | This work |
| pKM1 | pNTR-SD-*msrmsd* (Sen2) | pNTR-SD | Over-expression of *msrmsd* (Sen2). | Ampicillin (50 µg/mL) | This work |
| pJB137 | pNTR-SD-msrmsd(Eco9) | pNTR-SD | Over-expression of msrmsd(Eco9). | Ampicillin (50 µg/mL) | This work |
| pJB112 | pTU175-pBAD-msrmsd(Sen2)-rcaT(Eco9)-RT(Sen2) | pTU175 | Construction of plasmid pJB138. | Spectinomycin | This work |
| pJB130 | pTU175-msrmsd(Eco9)-rcaT(Sen2)-RT(Eco9) | pTU175 | Construction of plasmid pJB139. | Spectinomycin | This work |
| pJPS1 | pJPS1 | pJPS1 | Tagging *rrtT* with a 2x-Strep-3xFlag tag. | Ampicillin, Kanamycin | 53 |
| pKD4 | pKD4 | pKD4 | Gene deletion. | Ampicillin, Kanamycin | 36 |
| pKD45 | pKD45 | pKD45 | Scar-less *msd* deletions. | Ampicillin, Kanamycin | Datsenko & Wanner, unpublished |
| pKD46 | pKD46 | pKD46 | λ-red recombineering. | Ampicillin | 36 |
| pTU175 | pTU175-empty | pTU175 | Cloning. | Spectinomycin | 50 |
| pBAD30 | pBAD30-empty | pBAD30 | Cloning. | Ampicillin | 51 |
| pBAD33 | pBAD33-empty | pBAD33 | Cloning. | Chloramphenicol | 51 |
| pCP20 | pCP20 | pCP20 | Flipping-out resistance cassettes. | Ampicillin, Chloramphenicol | 38 |
| pET28α | pET28α-empty | pET28α | Cloning. | Kanamycin | Novagen |
| pNTR-SD | pNTR-SD-empty | pNT3 | Cloning. | Ampicillin (50 µg/mL) | 52 |
