## Supplementary Table 4. Description of construction of plasmids used in this study. for "Bacterial retrons encode tripartite toxin/antitoxin systems"

**Supplementary Table 4.** Description of construction of plasmids used in this study (1/2).

| Plasmid code | Plasmid construction comments | Vector | Insert |
|---|---|---|---|
| pJB11 | Insert was amplified using primers JB92 and JB93 from strain *S* Tm, and ligated in NheI-HindIII cut-pET28α. | pET28α | NheI-*rcaT*-HindIII |
| pJB12 | Insert was amplified using primers JB103 & JB93 from strain *S* Tm, and ligated in XmaI-HindIII cut-pBAD30. | pBAD30 | XmaI-*rcaT*-HindIII |
| pJB120 | Insert was amplified using primers JB419 & JB420 from plasmid pJB51, and ligated in NcoI-XhoI cut-pET28α. | pET28α | NcoI-*msrmsd*-FRT-*rrtT*-XhoI |
| pJB15 | Insert was amplified using primers JB103 & JB93 from strain *S* Tm, and ligated in XmaI-HindIII cut-pBAD33. | pBAD33 | XmaI-*rcaT*-HindIII |
| pJB24 | Insert was amplified using primers JB92 and JB93 from strain NT16150, and ligated in NheI-HindIII cut-pET28α. | pET28α | NheI-*rcaT*- D296V-HindIII |
| pJB37 | Insert was amplified using primers JB187 & JB93 from plasmid pJB15, and ligated in XhoI-HindIII cut-pTU175. | pTU175 | XhoI-pBAD-*rcaT*-HindIII |
| pJB38 | Insert was amplified using primers JB187 & JB99 from plasmid pJB16, and ligated in XhoI-HindIII cut-pTU175. | pTU175 | XhoI-pBAD-*msrmsd*-*rcaT*-*rrtT*-HindIII |
| pJB45 | Insert was amplified using primers JB101 & JB93 from strain *S* Tm, and ligated in XmaI-HindIII cut-pJB38. | pJB38 | XmaI-*msrmsd*-*rcaT*-HindIII |
| pJB46 | Insert was amplified using primers JB101 & JB93 from strain NT16074, and ligated in XmaI-HindIII cut-pJB38. | pJB38 | XmaI-*msr*Δ62*msd*-*rcaT*-HindIII |
| pJB47 | Insert was amplified using primers JB101 & JB93 from strain NT16489, and ligated in XmaI-HindIII cut-pJB38. | pJB38 | XmaI-*msr*Δ63*msd*-*rcaT*-HindIII |
| pJB48 | An *msd* region was deleted (Δ71:*msd*) by PCR-directed deletion using primers JB183 & JB184, based on plasmid-template pJB45. | pJB45 | XmaI-*msr*Δ71*msd*-*rcaT*-HindIII |
| pJB49 | An *msd* region was deleted (Δ79:*msd*) by PCR-directed deletion using primers JB183 & JB185, based on plasmid-template pJB45. | pJB45 | XmaI-*msr*Δ79*msd*-*rcaT*-HindIII |
| pJB51 | Insert was amplified using primers JB99 & JB101 from strain NT16277, and ligated in XmaI-HindIII cut-pJB38. | pJB38 | XmaI-*msrmsd*-FRT-*rrtT*-HindIII |
| pJB64 | An *msd* region was deleted (Δ67:*msd*) by PCR-directed deletion using primers JB183 & JB288, based on plasmid-template pJB45. | pJB45 | XmaI-*msr*Δ67*msd*-*rcaT*-HindIII |
| pJB86 | Insert was amplified using primers JB345 & JB99 from plasmid pJB38, and ligated in XmaI-HindIII cut-pJB38. | pJB38 | XmaI-branchingG→T*msrmsd*-*rcaT*-*rrtT*-HindIII |
| pJB87 | Insert was amplified using primers JB103 & JB99 from strain *S* Tm, and ligated in XmaI-HindIII cut-pJB38. | pJB38 | XmaI-*rcaT*-rrtT-HindIII |
| pJB91 | Insert was amplified using primers JB368 & JB369 from strain NT12549 (*E. coli* NILS-16), and ligated in XmaI-SalI cut-pBAD33. | pBAD33 | XmaI-rcaT(Eco9)-SalI |
| pJB92 | Insert was amplifed using primers JB370 & JB371 from strain NT12549 (*E. coli* NILS-16; NT12549), and ligated in XmaI-SalI cut-pBAD33. | pBAD33 | XmaI-msrmsd(Eco9)-rcaT(Eco9)-rt(Eco9)-SalI |
| pJB93 | Insert was amplified using primers JB370 & JB390 from plasmid pJB92, and ligated in XmaI-XbaI cut-pJB51. | pJB51 | XmaI-msrmsd(Eco9)-XbaI |
| pJB94 | Insert was amplified using primers JB391 & JB392 from plasmid pJB92, and ligated in XbaI-HindIII cut-pJB51. | pJB51 | XbaI-RT(Eco9)-HindIII |
| pJB95 | Insert was amplified using primers JB391 & JB392 from plasmid pJB92, and ligated in XbaI-HindIII cut-pJB93. | pJB93 | XbaI-RT(Eco9)-HindIII |
| pJB112 | Insert was amplified using primers JB443 & JB444 from plasmid pJB92, and ligated in XbaI cut-pJB51. | pJB51 | NheI-rcaT(Eco9)-NheI |

**Supplementary Table 4.** Description of construction of plasmids used in this study (2/2).

| | | | |
|---|---|---|---|
| pJB130 | Insert was amplified using primers JB445 & JB446 from plasmid pJB38, and ligated in XbaI cut-pJB95. | pJB95 | NheI-*rcaT* (Sen2)-NheI |
| pJB137 | Insert was amplified using primers JB455 & JB456 from plasmid pJB92, and ligated in HindIII-EcoRI cut-pNTR-SD. | pNTR-SD | HindIII-msrmsd(Eco9)-EcoRI |
| pJB138 | Insert was amplified using primers JB368 & JB99 from plasmid pJB112, and ligated in XmaI-HindIII cut-pJB38. | pJB38 | XmaI-rcaT(Eco9)-RT(Sen2)-HindIII |
| pJB139 | Insert was amplified using primers JB103 & JB392 from plasmid pJB130, and ligated in XmaI-HindIII cut-pJB38. | pJB38 | XmaI-rcaT(Sen2)-RT(Eco9)-HindIII |
| pJB140 | Insert was amplified using primers JB368 & JB392 from plasmid pJB92, and ligated in XmaI-HindIII cut-pJB38. | pJB38 | XmaI-rcaT(Eco9)-RT(Eco9)-HindIII |
| pJB141 | Insert was amplified using primers JB457 & JB99 from plasmid pJB38, and ligated in XmaI-HindIII cut-pJB38. | pJB38 | XmaI-SD-RT(Sen2)-HindIII |
| pJB142 | Insert was amplified using primers JB458 & JB392 from plasmid pJB92, and ligated in XmaI-HindIII cut-pJB38. | pJB38 | XmaI-SD-RT(Eco9)-HindIII |
| pKM1 | Insert was amplified using primers KM_R_01 & KM_R_03 from plasmid pJB38, and ligated in HindIII-SalI cut-pNTR-SD. | pNTR-SD | HindIII-*msrmsd* (Sen2)-SalI |
