## Supplementary Table 5. List of primers used in this study. for "Bacterial retrons encode tripartite toxin/antitoxin systems"

**Supplementary Table 5.** List of primers used in this study (1/2).

| Primer code | Primer name | Primer sequence | Primer description | Used in |
|---|---|---|---|---|
| JB92 | rcaT_NheI_For | TTCATCATGCTAGC ATGTTTGAAGAAACTATTAAAAAACAATTT | Green: anneals to start of *rcaT*, blue: restriction site, black: restriction enzyme buffer sequence. | pJB11, pJB24 plasmid construction. |
| JB93 | rcaT_HindIII_Rev | TTCATCATAAGCTT TTAATTAATCTCCTCAGCCATT | Green: anneals to end of *rcaT*, blue: restriction site, black: restriction enzyme buffer sequence. | pJB11, pJB12, pJB15, pJB24, pJB37, pJB45, pJB46, pJB47 plasmid construction. |
| JB103 | rcaT_XmaI_For | TTCATCATCCCGGG ATGTTTGAAGAAACTATTAAAAAACAATTT | Green: anneals to start of *rcaT*, blue: restriction site, black: restriction enzyme buffer sequence. | pJB12, pJB15, pJB87, pJB139 plasmid construction. |
| JB101 | msr_XmaI_For | TTCATCATCCCGGG ACATCACTCTTTAGCGTTAG | Green: anneals to start of *msr*-Sen2, blue: restriction site, black: restriction enzyme buffer sequence. | pJB45, pJB46, pJB47, pJB51 plasmid construction. |
| JB187 | pBAD_XhoI_For | TTCATCATCTCGAG GCATAATGTGCCTGTCAAATGGAC | Green: anneals to start of pBAD promoter, blue: restriction site, black: restriction enzyme buffer sequence. | pJB37, pJB38 plasmid construction. |
| JB99 | rrtT_HindIII_Rev | TTCATCATAAGCTT TTATTTGTCTTTTTGTTTTTG | Green: anneals to end of *rrtT*, blue: restriction site, black: restriction enzyme buffer sequence. | pJB38, pJB51, pJB87, pJB138, pJB141 plasmid construction. |
| JB345 | msr-branchingGtoT_XmaI_For | TTCATCATCCCGGG ACATCACTCTTTATCGTTAG | Green: anneals to start of *msd*, red: branching G to T point mutation, blue: restriction site, black: restriction enzyme buffer sequence. | pJB86 plasmid construction. |
| JB368 | rcaT(Eco9)_XmaI_For | TTCATCATCCCGGG atgAGTGATTCTGCATATATGAATGCATTAC | Green: anneals to start of rcaT-Eco9, blue: restriction site, black: restriction enzyme buffer sequence. | pJB91, pJB138, pJB140 plasmid construction. |
| JB369 | rcaT(Eco9)_SalI_Rev | TTCATCATGTCGAC CcatTctaAGCCCCTCCA | Green: anneals to end of rcaT-Eco9, blue: restriction site, black: restriction enzyme buffer sequence. | pJB91 plasmid construction. |
| JB370 | msr(Eco9)_XmaI_For | TTCATCATCCCGGG ATCTCGCATCTTTTGCTATCCTAACC | Green: anneals to start of msr-Eco9, blue: restriction site, black: restriction enzyme buffer sequence. | pJB92, pJB93 plasmid construction. |
| JB371 | RT(Eco9)_SalI_Rev | TTCATCATGTCGAC GCGCAAGCTAGATAAAGTCATGCT | Green: anneals to end of rt-Eco9, blue: restriction site, black: restriction enzyme buffer sequence. | pJB92 plasmid construction. |
| JB390 | msd(Eco9)_XbaI_Rev | TTCATCATTCTAGA CCACCACAAACAAAAATAATAGGAGGCAT | Green: anneals to +107 nts of rcaT-Eco9, blue: restriction site, black: restriction enzyme buffer sequence. | pJB93 plasmid construction. |
| JB391 | RT(Eco9)_XbaI_For | TTCATCATTCTAGA atgGAGTTGCTAAGTTATATTTCAAAAGGTC | Green: anneals to start of rt-Eco9, blue: restriction site, black: restriction enzyme buffer sequence. | pJB94, pJB95 plasmid construction. |
| JB392 | RT(Eco9)_HindIII_Rev | TTCATCATAAGCTT ttaAATTTCATATTTTGAAATTTCGCTGATTACATCATCC | Green: anneals to end of rt-Eco9, blue: restriction site, black: restriction enzyme buffer sequence. | pJB94, pJB95, pJB139, pJB140, pJB142 plasmid construction. |
| JB419 | msr_NcoI_For | TTCATCATCCATGG ACATCACTCTTTAGCGTTAG | Green: anneals to start of *msr*-Sen2, blue: restriction site, black: restriction enzyme buffer sequence. | pJB120 plasmid construction. |
| JB420 | rrtT-TEV_XhoI_Rev | TTCATCATCTCGAG tgcattggattggaagtacaggttttcctcgat aga gcc gcc gcc gcc TTTGTCTTTTTGTTTTTGT | Green: anneals to end of *rrtT*, purple: linker, orange: TEV-protease site, blue: restriction site, black: restriction enzyme buffer sequence. | pJB120 plasmid construction. |
| JB183 | msrmsd_Rev | AGTACTCAATAAGTTAGTGTCTGGCGAAAC | Green: anneals close to start of *msr.* | pJB48, pJB49, pJB64 plasmid construction. |
| JB184 | Δ71msd_For | CTCCTCTAGTAAAGAGTAAAAAACGAGC | Green: anneals to *msd.* | pJB48 plasmid construction. |
| JB185 | Δ79msd_For | GTAAAGAGTAAAAAACGAGCAAAAAACATTGC | Green: anneals to *msd.* | pJB49 plasmid construction. |
| JB288 | Δ67msd_For | GCGCCTCCTCTAGTAAAGAGTAAAAAAC | Green: anneals to *msd.* | pJB64 plasmid construction. |
| JB443 | rcaT-Eco9_NheI_For | TTCATCATGCTAGC atgAGTGATTCTGCATATATGAATGCATTAC | Green: anneals to start of rcaT-Eco9, blue: restriction site, black: restriction enzyme buffer sequence. | pJB112 plasmid construction. |
| JB444 | rcaT-Eco9_NheI_Rev | TTCATCATGCTAGC CcatTctaAGCCCCTCCA | Green: anneals to end of rcaT-Eco9, blue: restriction site, black: restriction enzyme buffer sequence. | pJB112 plasmid construction. |
| JB445 | rcaT-Sen2_NheI_For | TTCATCATGCTAGC AAAAAACGAGCAAAAAACATTGCTCG | Green: anneals 75 nucleotides upstream of *rcaT*-Sen2 start, blue: restriction site, black: restriction enzyme buffer sequence. | pJB130 plasmid construction. |
| JB446 | rcaT-Sen2_NheI_Rev | TTCATCATGCTAGC CTTTCGAAGGATGAGCAATAGTTCTC | Green: anneals 150 nucleotides downstream of *rcaT*-Sen2 end, blue: restriction site, black: restriction enzyme buffer sequence. | pJB130 plasmid construction. |
| JB455 | msrmsd-Eco9_HindIII_For | TTCATCATAAGCTT CCTACCTTTACGCGCG | Green: anneals to start of msr-Eco9, blue: restriction site, black: restriction enzyme buffer sequence. | pJB137 plasmid construction. |
| JB456 | msrmsd-Eco9_EcoRI_Rev | TTCATCATGAATTC GTGACCCCCTACTCCA | Green: anneals 49 nucleotides downstream of msd-Eco9 end, blue: restriction site, black: restriction enzyme buffer sequence. | pJB137 plasmid construction. |
| JB457 | RT-Sen2_SD_XmaI_For | TTCATCATCCCGGG aggaggAAT atgGACATATTACAGCATATTTC | Green: anneals to start of *rrtT*, orange: Shine-Dalgarno site, blue: restriction site, black: restriction enzyme buffer sequence. | pJB141 plasmid construction |
| JB458 | RT-Eco9_SD_XmaI_For | TTCATCATCCCGGG aggaggAAT atgGAGTTGCTAAGTTATATTTCAAAAGGTC | Green: anneals to start of rt-Eco9, orange: Shine-Dalgarno site, blue: restriction site, black: restriction enzyme buffer sequence. | pJB142 plasmid construction |
| KM_R_01 | msrmsd-Sen2_For | CAAGCAAGCTT ACATCACTCTTTAGCGTTAGG | Green: anneals to start of *msr*-Sen2, blue: restriction site, black: restriction enzyme buffer sequence. | pKM1 plasmid construction. |
| KM_R_03 | msrmsd-Sen2_Rev | GAGCTGTCGAC CAAACATTGATTTAAACGTTATG | Green: anneals 82 nucleotides downstream of *msd*-Sen2 end, blue: restriction site, black: restriction enzyme buffer sequence. | pKM1 plasmid construction. |
| JB464 | rrtT_Cter-pJPS1_For | TTAAATATGGATCTGATAATATTTATAAATTACAAAAACAAAAAGACAAA GGCTGGTCACACCCGCAGTTTG | Green: anneals to C-terminus of *rrtT*, blue: anneals to plasmid pJPS1 (to amplify 2xStrep-TEV-3xFlag:kan). | Forward primer to amplify cassette for chromosomal *rrtT*-2xStrep-3xFlag tagging. |
| JB465 | rrtT_Cter-pJPS1_Rev | GGAAAACTTATTCCAAGTTGTTCTGGGCATAATCAACAAAACACTGGTGT CATATGAATATCCTCCTTAG | Green: anneals down-stream of C-terminus of *rrtT*, blue: anneals to plasmid pJPS1 (to amplify 2xStrep-TEV-3xFlag:kan). | Reverse primer to amplify cassette for chromosomal *rrtT*-2xStrep-3xFlag tagging. |
| JB319 | rcaT_Cter_pKD45_For | CAGCGCTGCGTCAAATAATAGCAAAAAAGAAATGGCTGAGGAGATTAAT CGCATCGTGGCCGGATCTTGC | Green: anneals to C-terminus of *rcaT*, blue: anneals to plasmid pKD45 (to amplify PrhaBAD-*ccdB:kan*). | Forward primer to amplify cassette for chromosomal insertion of PrhaBAD-*ccdB*-kan at the *rcaT*-C-terminus. |
| JB320 | rcaT_Cter_pKD45_Rev | AAAGGAAATGATTTCACTTTTCTTGGTTAATAGTAAATCTGAAATATGCT CGGATAACAGAAAGGCCGGG | Green: anneals down-stream of C-terminus of *rcaT*, blue: anneals to plasmid pKD45 (to amplify PrhaBAD-*ccdB:kan*). | Reverse primer to amplify cassette for chromosomal insertion of PrhaBAD-*ccdB:*kan at the *rcaT*-C-terminus. |
| JB321 | rcaT-3xFlag_Cter_For | CAGCGCTGCGTCAAATAATAGCAAAAAAGAAATGGCTGAGGAGATTAAT ggc ggc ggc ggc tct GACTACAAAGACCATG | Green: anneals to C-terminus of *rcaT*, blue: anneals to plasmid pJPS1 (to amplify 3xFlag), orange: linker. | Forward primer to amplify fragment for chromosomal *rcaT*-3xFlag tagging. |
| JB322 | rcaT-3xFlag_Cter_Rev | AAAGGAAATGATTTCACTTTTCTTGGTTAATAGTAAATCTGAAATATGCTGTAATAT GTCcatta CTTGTCATCGTCATCC | Green: anneals to C-terminus of *rcaT*, blue: anneals to plasmid pJPS1 (to amplify 3xFlag). | Reverse primer to amplify fragment for chromosomal *rcaT*-3xFlag tagging. |

**Supplementary Table 5.** List of primers used in this study (2/2).

| | | | | |
|---|---|---|---|---|
| JB348 | Δ79_msd_pKD45_F | GATTTATAGCCTTGTCGAGCGTTTCGCCAGACACTAACTTATTGAGTACT CGCATCGTGGCCGGATCTTGC | Green: anneals to *msr* region, blue: anneals to plasmid pKD45 (to amplify PrhaBAD-*ccdB:kan*). | Forward primer to amplify cassette for chromosomal insertion of a Δ79:*msd*-PrhaBAD-*ccdB*-kan, replacing the *msrmsd* locus. |
| JB349 | Δ79_msd_pKD45_R | CGTTGATGAAAAAAACGAGCAATGTTTTTTGCTCGTTTTTTACTCTTTAC CGGATAACAGAAAGGCCGGG | Green: anneals to end of *msd* region, blue: anneals to plasmid pKD45 (to amplify PrhaBAD-*ccdB:kan*). | Reverse primer to amplify cassette for chromosomal insertion of a Δ79:*msd*-PrhaBAD-*ccdB*-kan, replacing the *msrmsd* locus. |
| JB350 | msr_For | ACATCACTCTTTAGCGTTAGGCTTTGATTTATAGCCTTGTCGAGCGTTTC | Green: anneals to *msr* region. | Forward primer to amplify fragment for chromosomal *msd* deletions (fragments amplified from pJB45, pJB46, pJB47, pJB48, pJB49, or pJB64). |
| JB351 | rcaT-internal_Rev | GCTGTGTAAGTGAGTAGTCTATCCCTAAAACTGGGGGGAATTGGTGCACG | Green: anneals to internal region of *rcaT*. | Reverse primer to amplify fragment for chromosomal *msd* deletions (fragments amplified from pJB45, pJB46, pJB47, pJB48, pJB49, or pJB64). |
| JB104 | araBAD_pKD13_Del_F | ACTCTCTACTGTTTCTCCATACCTGTTTTTCTGGATGGAGTAAGACGATG ATTCCGGGGATCCGTCGACC | Green: anneals to start of *S*Tm *araBAD* operon, black: anneals to plasmid pKD13. | Deletion of *araBAD* operon in *S*Tm. |
| JB105 | araBAD_pKD13_Del_R | GCCCCCCATGGGACGCGTTTTTAGAGGCATTACTGCCCGTAATAGGCTTT GTAGGCTGGAGCTGCTTCG | Green: anneals to end of *S*Tm *araBAD* operon, black: anneals to plasmid pKD13. | Deletion of *araBAD* operon in *S*Tm. |
| JB84 | rnhA_pKD4_Del_F | AGACTTGCGTTTTGTTATCGATTTCAATTACAGGAAGTCTACCAGAGATG GTGTAGGCTGGAGCTGCTTC | Green: anneals to start of *S*Tm *rnhA*, black: anneals to plasmid pKD4. | Deletion of *rnhA* in *S*Tm (NT16276). |
| JB85 | rnhA_pKD4_Del_R | GCCACCCGGCAATATCGCAAACCGGATGGCTAAGCTTCCGCCTGGTAGCC CATATGAATATCCTCCTTA | Green: anneals to end of *S*Tm *rnhA*, black: anneals to plasmid pKD4. | Deletion of *rnhA* in *S*Tm (NT16276). |
| JB172 | xseB_pkD4_F | CGTTACCATAGACTTGTACTGCCATTACAGAGAGTCAAACAGTCATCATG GTGTAGGCTGGAGCTGCTTC | Green: anneals to start of *S*Tm *xseB*, black: anneals to plasmid pKD4. | Deletion of *xseB* in *S*Tm (NT16487). |
| JB173 | xseB_pkD4_R | CAAGCCTGTAGCTGCTGCGTAAAGTCCATTTACTCATTATCCGCGATAAA CATATGAATATCCTCCTTA | Green: anneals to end of STm xseB, black: anneals to plasmid pKD4. | Deletion of *xseB* in *S*Tm (NT16487). |
| JRE_Δmsrmsd1 | msrA_For | ACCGGTTGAAAAAGATGCACTTCTCAACCATAATAACATC GTGCAGGCTGGAGCTGCTTC | Green: anneals to start of *msr*-Sen2, black: anneals to plasmid pKD4. | Deletion of *msrmsd*-Sen2 in *S*Tm (NT16085). |
| JRE_Δmsrmsd2 | msdB_Rev | TGTTTTTTGCTCGTTTTTTACTCTTTACTAGAGGAGGCGC CATATGAATATCCTCCTTAG | Green: anneals to end of *msd*-Sen2, black: anneals to plasmid pKD4. | Deletion of *msrmsd*-Sen2 in *S*Tm (NT16085). |
